# Molecular Neurobiology of Loss

**DOI:** 10.1101/2022.07.13.499899

**Authors:** Marissa A. Smail, Brittany L. Smith, Rammohan Shukla, Khaled Alganem, Hunter M. Eby, Justin L. Bollinger, Ria K. Parikh, James B. Chambers, James K. Reigle, Rachel D. Moloney, Nawshaba Nawreen, Eric S. Wohleb, Harry Pantazopoulos, Robert E. McCullumsmith, James P. Herman

## Abstract

Psychological loss is a common experience that erodes well-being and negatively impacts quality of life. The molecular underpinnings of loss are poorly understood. Here, we investigate the mechanisms of loss using an enrichment removal (ER) paradigm in rats. A comprehensive multi-omics investigation of the basolateral amygdala (BLA) revealed alterations in microglia and extracellular matrix (ECM). Follow-up studies indicated that ER decreased microglia size, complexity, and phagocytosis, suggesting reduced immune surveillance. Loss also substantially increased ECM coverage, specifically targeting perineuronal nets surrounding parvalbumin interneurons, suggesting decreased plasticity and increased inhibition in the BLA following loss. Behavioral analyses suggest that these molecular effects are linked to impaired BLA salience evaluation, reflecting emotional blunting observed in human loss. These loss-like behaviors could be rescued by depleting BLA ECM during removal, helping us understand the mechanisms underlying loss and revealing novel molecular targets to ameliorate its impact.

## INTRODUCTION

Psychological loss is something that most people will experience in their lifetime. Losing something of value (e.g., interpersonal relationships, financial stability, secure housing, health) can erode well-being and negatively impact quality of life^1–3^. The impact of loss is felt in particular during times of natural disasters, political upheavals, and pandemics^4^.

Loss can be defined as a “state of deprivation of a motivationally significant conspecific, object, or situation^1^.” This relative deprivation often precipitates a depressed emotional state, and symptoms of loss can resemble those seen in major depressive disorder (MDD), including amotivation, sadness, withdrawal, rumination, and compromised executive function^1–3^. Loss is also associated with ‘atypical’ MDD symptoms such as weight gain and hypoactive hypothalamic-pituitary-adrenal (HPA) axis responsivity^5,6^. Little is known about the molecular mechanisms underlying this experience, as the unique phenotype is difficult to track clinically and has received little attention in preclinical studies^5–7^. Elucidating the molecular mechanisms driving the response to loss could reveal novel treatments that may be of widespread benefit.

Our lab recently developed a model of loss in rats, in which environmental enrichment (EE) serves as proxy for positive life events and its removal emulates loss^8,9^. EE is known to have rewarding properties in rodents, providing social, physical, and cognitive stimulation that contribute to enhanced cognition and neuroplasticity, as well as beneficial effects on emotional reactivity to stress^10,11^. The enrichment removal (ER) protocol begins with 4 weeks of EE followed by removal to single housing lacking availability of positive stimuli. This negative transition from high to low environmental stimulation is consistent with removal of a motivationally significant condition. Indeed, after 1 week of ER, adult male rats exhibit “loss-like” phenotypes, including increased passive coping in the forced swim test, increased hedonic drive in the sucrose preference test, weight gain, and a hypoactive HPA axis relative to EE and standard housed (SH) rats^8^. Thus, ER affords a unique opportunity to study loss-related pathologies in a rodent model.

Here, we utilize a series of complementary approaches to investigate the biological substrates of psychological loss. Post-behavior (forced swim) Fos expression revealed selective recruitment of BLA neurons following ER. We then used a discovery-based multi-omics approach to build molecular signatures of ER in the BLA. Bioinformatics analyses of these signatures identified microglia and the extracellular matrix as consistently altered targets. Follow up molecular and behavioral studies revealed how ER-induced changes in these candidate mechanisms impact BLA plasticity and BLA-dependent behaviors. We then demonstrated that these loss-like phenotypes could be rescued by enzymatically depleting the BLA extracellular matrix during the removal period. Taken together, this information-rich dataset provides unique insight into a role of the extracellular matrix in driving diverse neural and behavioral dysfunctions consequent to loss in the amygdala.

## RESULTS

### Experiment 1: BLA Exhibits Differential Fos Activation Following ER

We initially ran a post-forced swim Fos screen to identify regions that may be contributing to loss phenotypes (Figure 1A), focusing on known stress-responsive brain regions. The most notable pattern of Fos expression was observed in the basolateral amygdala (BLA), where ER produced a pronounced increase in the number of Fos reactive neurons relative to continuously-enriched (EE) controls (Figure 1B; Extended Data 2). Given the established role of the BLA in stress processing and adaptation^12,13^, we chose to focus on this region for further analysis (Table S1).

**Figure 1:**
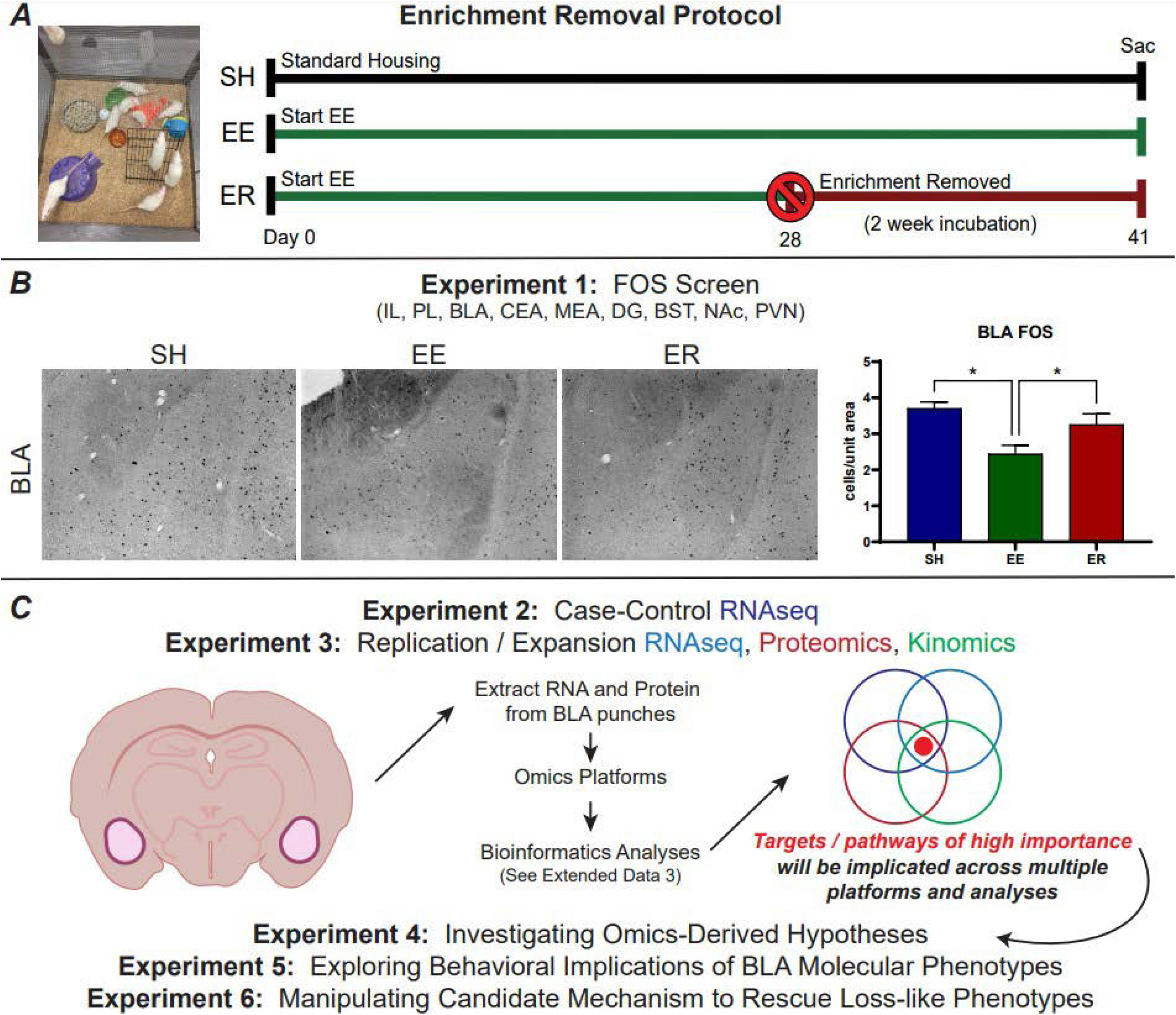
Experimental Background and Overview. (A) Example of environmental enrichment (EE), and timeline for standard enrichment removal (ER) protocol. (B) Representative images of the BLA from initial Fos screen and quantification (n=8-9/group). (C) Outline for omics and follow-up studies. Experiments 2 and 3 utilized multi-omics in tandem with a number of complementary bioinformatics analyses to explore the molecular landscape of the BLA following ER. Experiments 4 and 5 further explored the hypotheses that emerged from these analyses, as well as their role in ER behavioral phenotypes, to expand our understanding of novel mechanisms that underlie loss. Experiment 6 explores manipulating the molecular mechanisms identified here in an attempt to rescue the loss-like behavioral phenotypes generated by ER. * = p<0.05.

### Experiment 2: RNAseq Points to Immune and Matrix Alterations Following ER

We then designed a series of hypothesis-generating experiments to uncover potential mechanisms underlying the effects of ER in the BLA. This consisted of case-control RNAseq run on BLA micropunches from SH, EE, and ER rats (Figure 1C). Analyzing data as complex as transcriptomic signatures with a singular platform is inherently biased and could easily miss novel information that could be gleaned from broader, more comprehensive analyses encompassing multiple levels and perspectives. Here we leveraged a pipeline that covers multiple dimensions of transcriptomic analyses and thus lends more confidence to interpretation of gene expression patterns (Extended Data 3).

In a second cohort of rats, micropunches of the BLA were collected^14^, RNA was extracted and samples were submitted for RNAseq. Transcriptomic data was generated for SH, EE, and ER rats, yielding signatures for 3 contrasts: EEvSH, ERvSH and ERvEE (Table S2). EEvSH represents changes generated by the experience of enrichment, ERvSH represents changes generated by the full experience of enrichment and removal, and ERvEE represents changes generated specifically by removal. Both ERvSH and ERvEE contrasts are important to understanding ER phenotypes, as comparison to EE (procedural control group) indicates how loss impacts the BLA and comparison to SH (baseline control group) indicates how those changes compare to rats that have only experienced unstimulated conditions.

#### Full Transcriptome Pathway Analysis

The first analysis used gene set enrichment analysis (GSEA)^15^ to broadly assess gene expression across the full transcriptome in the BLA. Significantly enriched gene ontology pathways were identified for each contrast and organized into different functional categories using PathwayHunter^16^ (see Methods; Figure 2A left, small labels; Extended Data 4A; Table S4). These categories were further condensed based on *a priori* knowledge (Figure 2A left, large labels), and the strength and direction of changes in these categories was determined based on hypergeometric overlap (Fig 2A left, -log10 p values) and GSEA enrichment score (Figure 2A left, colors), respectively. Many of the observed pathways were upregulated in EE (EEvSH, yellow) and downregulated in ER (ERvEE, blue). Strongly implicated functional categories included *immune, extracellular signaling, development, regulation, response to stimulus*, and *cell structure*.

**Figure 2:**
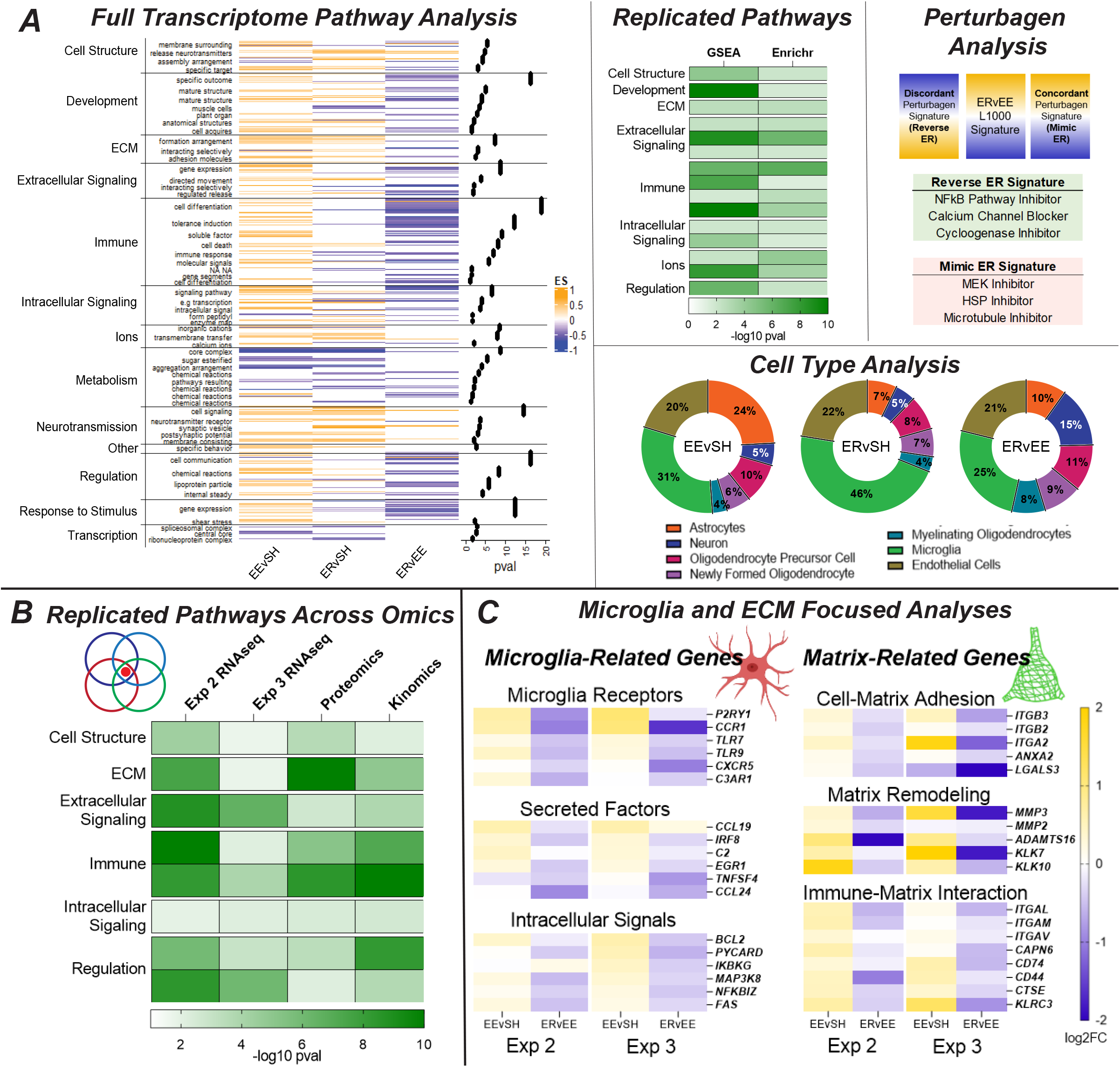
Multi-Omics Indicates ER Alterations to Microglia and the Extracellular Matrix. (A) Results from Experiment 2 (n=6/group) (Table S4). The first analysis utilized GSEA to conduct full transcriptome pathway analysis. Significant pathways (p<0.05) were condensed based on semantic similarity using Pathway Hunter (small left labels, and these categories were further condensed into functional themes based on *a priori* knowledge (leftmost labels)). Enrichment scores for individual pathways are represented by the heatmap (with EEvSH on the left, ERvSH in the middle, and ERvEE on the right), and the degree of each category’s enrichment is represented by the p value dots on the right. The enrichment scores show how much (magnitude) and in what direction (yellow is up, blue is down) a theme was changed by the model, while the p values show the contribution of each category to these changes. The second analysis utilized Enrichr to conduct targeted pathway analysis and generated a similar set of results (Extended Data 4B). Themes that replicated between these two pathway analyses are shown in the replicated pathways heatmap, with darker green indicating more involvement. The third analysis utilized iLINCS to look for compounds that would replicate (concordant) or reverse (discordant) the present phenotypes (Table S5). We also used the leading-edge genes from GSEA to conduct cell type analysis (Table S6). Collectively, these pipeline results start to implicate microglia and the extracellular matrix (ECM) in ER. (B) Similar analyses were run on the data from Experiment 3 (n=10/group pooled). The RNAseq results are presented in Extended Data 5, while the proteomics and kinomics results are presented in Extended Data 6. Those results are summarized and compared to the above results here, and they provide more support for microglia and the ECM. This replication across cohorts, omics platforms, and analyses gives us higher confidence in these targets’ involvement. (C) Hypothesis-driven analyses of microglia and ECM-related genes further implicates both at multiple levels and points to a pattern of upregulation in EE (yellow), followed by downregulation in ER (blue) (Table S7).

#### Targeted Pathway Analysis

The second analysis focused on the strongest differential expression. The top 100 upregulated and top 100 downregulated genes (ranked by highest fold change) were determined for each contrast and entered into Enrichr for pathway analysis^17^. The results were condensed in the same manner as the full transcriptome pathways and yielded a similar pattern of upregulation in EE and downregulation in ER (Extended Data 4B; Table S4). While fewer overall pathways were found in this dataset, we again observed strong involvement of *immune, extracellular signaling*, and *regulation* categories.

Directly comparing the functional categories implicated by both full transcriptome and targeted pathway analysis emphasizes the consistencies between the two approaches (Figure 2A middle, Extended Data 4C; Table S4). Specifically, themes observed in both analyses were: *extracellular matrix, extracellular signaling, immune, cell structure, development, intracellular signaling, ions*, and *regulation*.

#### Signature Analysis

Our third analysis utilized iLINCS (integrated network-based cellular signatures), a publicly available database curating transcriptomic signatures for thousands of drugs and compounds (aka perturbagens)^18^. iLINCS’s signatures are based on the L1000, a set of 978 genes that were measured in cell lines treated with these perturbagens. L1000 signatures were extracted for each contrast and uploaded to iLINCS. Similar perturbagen signatures were “concordant,” meaning that the compounds would be expected to recapitulate the experimental phenotype. On the other hand, different perturbagen signatures were “discordant,” meaning that the compounds would be expected to reverse (aka treat) the experimental phenotype (Figure 2A, top right). We primarily use these data to support the pathway analyses, as cell line signatures may not be fully comparable to the present bulk tissue micropunch data.

Findings here lend additional support for our pathway results. Discordant perturbagens involved the immune system (*NFkB and cyclooxygenase inhibitors*) and signaling (*calcium channel blocker*), while concordant perturbagens involved cell structure (*microtubule inhibitor*), response to stimulus (*HSP inhibitor*), and signaling *(MEK inhibitor*) (Figure 2A, top right). Looking more broadly at mechanism of action (MOA) across the different contrasts revealed that the strongest discordant perturbagens would impact the immune system, while the strongest concordant perturbagens would impact kinase signaling, suggesting specific signaling mechanisms that may be involved in loss (Extended Data 4D; Table S5).

#### Cell Type Analysis

We then used the leading-edge genes obtained from GSEA (i.e., the genes that drove the enrichment of significant pathways), to determine the contribution of individual cell types to the pathway results. The top 100 upregulated and top 100 downregulated leading-edge genes (by number of pathways) were uploaded into Kaleidoscope and their typical cell-type enrichment was determined using the “Brain RNA-Seq” module^19^. Proportions of this enrichment were determined to reveal the relative contribution of different cell types. Microglia exhibited the biggest contribution across all contrasts (Figure 2A, right middle; Table S6), followed by endothelial cells, astrocytes, and oligodendrocytes. Neuronal involvement was relatively small, supporting the notion that loss mechanisms favor the brain’s support systems over direct effects on neuronal circuitry itself.

### Experiment 3: Multi-Omics Supports Microglia and Extracellular Matrix Phenotypes

We next sought to validate and expand our ER signatures in a new cohort. This experiment provided 1) replication RNAseq and 2) omics data from other platforms that could potentially reveal whether changes observed at the RNA level survived to the protein (shotgun proteomics) and activity (serine-threonine kinomics) levels. BLA micropunches from a new cohort were subjected to our “triple prep” protocol (see Methods) allowing for simultaneous RNA and protein extraction from the same tissue. This preparation ensures that changes observed on multiple platforms were real and not an artifact of tissue heterogeneity. Extracted samples were then processed for parallel RNAseq, proteomics, and kinomics. Samples were pooled within groups.

#### RNAseq Analysis

The RNAseq was analyzed in the same manner as the previous cohort (Table S2). While individual pathways showed some variation (Extended Data 5A,B; Table S4), the functional categories associated with ER were similar to those observed in Experiment 2 (Extended Data 5C; Table S4). Similar patterns of upregulation in EE and downregulation in ER were observed in this cohort, as well as strong involvement of the immune system, extracellular matrix (ECM), and regulation across full and targeted pathway analyses. Signature analysis again implicated kinases and the immune system (Extended Data 5D; Table S5), while cell type analysis again pointed to microglia as the most involved cell type (Extended Data 5E; Table S6).

#### Proteomics and Kinomics Analysis

This experiment also offered insight into ER-related changes in protein and activity. EEvSH, ERvSH, and ERvEE signatures were generated for each platform (Table S3), and targeted pathway analysis was run in Enrichr. The targeted method was selected over the full transcriptome method because neither of these platforms yields data from enough targets to properly conduct the latter. Proteomics signatures were based on the top 100 up- and down-regulated peptides (based on fold change), while kinomics signatures were based on differentially phosphorylated peptides (FC>1.2).

The proteomics exhibited more upregulation across the different contrasts but involved similar themes as the RNAseq, including strong involvement of the ECM, immune system, and cell structure (Extended Data 6A; Table S4). The kinomics yielded a pattern similar to that of the gene expression data, with upregulation in EE followed by downregulation in ER, and implicated the immune system, cell cycle, regulation, and signaling (Extended Data 6B; Table S4). Upstream kinase analysis was also performed using KRSA, pointing to several kinases that may be implicated in ER (Extended Data 6C; Table S3)^20^.

While specific signatures from each omics platform varied to a degree, commonly implicated RNAseq themes were largely supported by both the proteomics and kinomics (Extended Data 6D; Table S4). Both indicated involvement of the immune system and ECM, indicating that these changes survive across multiple molecular domains of the BLA following ER. In combination, these studies give us high confidence that the experience of loss involves changes in BLA microglia and ECM (Figure 2B; Table S4).

### Hypothesis-Guided Analysis of Microglia and ECM Genes

Given the emergent roles of microglia and ECM in our hypothesis-free analyses, we went back into the omics datasets with this notion in mind. Looking specifically at genes associated with microglia, ECM, and the interaction of the two^21–23^, we observed strong patterns of upregulation in EE and downregulation in ER, suggesting a loss of function with microglia activity and ECM regulation following ER (Figure 2C; Table S7). Genes related to microglia receptors, secreted factors, and intracellular signals suggest that microglia homeostatic functions related to surveying and interacting with their surroundings may be attenuated^21,24–26^. Genes related to cell-matrix adhesion and matrix remodeling suggest that the ECM may not be assembling or degrading in the expected manner^27–29^. Genes related to immune-matrix interaction suggest that these alterations could be linked^22,30,31^.

### Experiment 4: Validating and Expanding Omics Hypotheses

A new cohort of ER rats was generated to further explore these mechanisms. For this cohort, rats were perfused, and immunohistochemistry (IHC) was used to validate and elaborate upon microglia and ECM phenotypes.

#### BLA Microglia Are Smaller, Less Complex, and Less Phagocytic Following ER

IBA1 (microglial marker) staining revealed that IBA1-immunoreactive area/microglia was reduced in both EE and ER relative to SH in the BLA (Figure 3A). However, we did not observe changes in microglia counts or intensity between groups (Extended Data 7A,B). To further dissect EE and ER phenotypes, we carried out a more detailed analysis of microglia morphology. Microglia soma circumference was not different between groups (Extended Data 7C). We then used the filament tracing tool in Imaris to reconstruct and analyze microglial processes from our 3 groups. The ER group alone exhibited shorter and smaller processes, fewer branch points, and fewer terminal points (Figure 3B). We then tested microglia function by co-staining with CD68, a lysosome marker associated with phagocytosis. A robust decrease of CD68 area/microglia was observed in the ER group, suggesting that ER microglia are less phagocytic (Figure 3C). These results align well with the omics that suggested ER microglia exhibit less surveillance of and interaction with their surroundings (Figure 3D)^21,25,26^.

**Figure 3:**
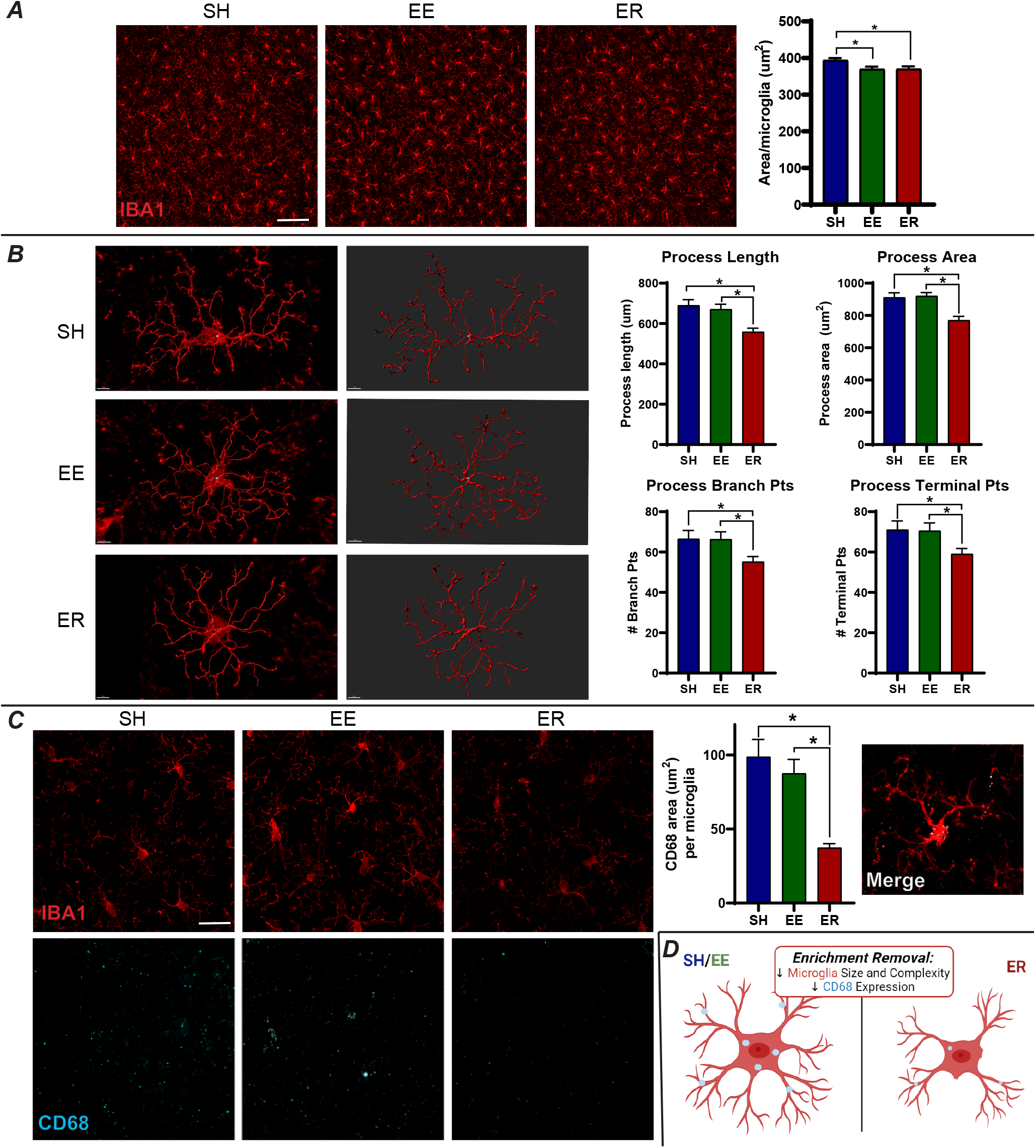
Microglia are Smaller, Less Complex, and Less Phagocytic Following ER. (A) Initial analysis of BLA microglia (labeled with IBA1) suggested that microglia are smaller in EE and ER (n=10/group; scale bar = 100 μm). (B) A more robust analysis of microglia morphology was conducted using the filament tracer tool in Imaris (n=8 microglia/animal) and revealed that, in ER alone, microglia are smaller and less complex. (C) ER rats express significantly less CD68 than SH or EE rats, suggesting that they are less phagocytic (scale bar = 10 μm). Merge shows localization of CD68 to microglia. (D) Summary of ER effects on microglia phenotypes. * = p<0.05.

#### ER Increases BLA Extracellular Matrix and Perineuronal Nets

The other main multi-omics finding was decreased regulation of ECM structure in ER. We used histochemistry to assess BLA expression of the ECM marker WFA (Wisteria Floribunda Agglutinin), which labels CSPGs (Chondroitin Sulfate Proteoglycans) of the ECM^27,28^. CSPGs are a critical component of perineuronal nets (PNNs), mesh-like ECM formations that surround neurons and help to regulate cell physiology. PNNs play a large role in neuroplasticity^29,32,33^, making them a potentially intriguing target in ER. As PNNs primarily surround inhibitory parvalbumin (PV) interneurons^27,28^, we co-labeled tissue for PV. WFA staining area was increased in ER rats relative to both EE and SH controls, as was the ratio of PV-immunoreactive cells decorated with PNNs (Figure 4A-C). WFA intensity followed a similar pattern of increasing in ER (Extended Data 7D), while the number of PV cells did not change between groups (Extended Data 7E).

**Figure 4:**
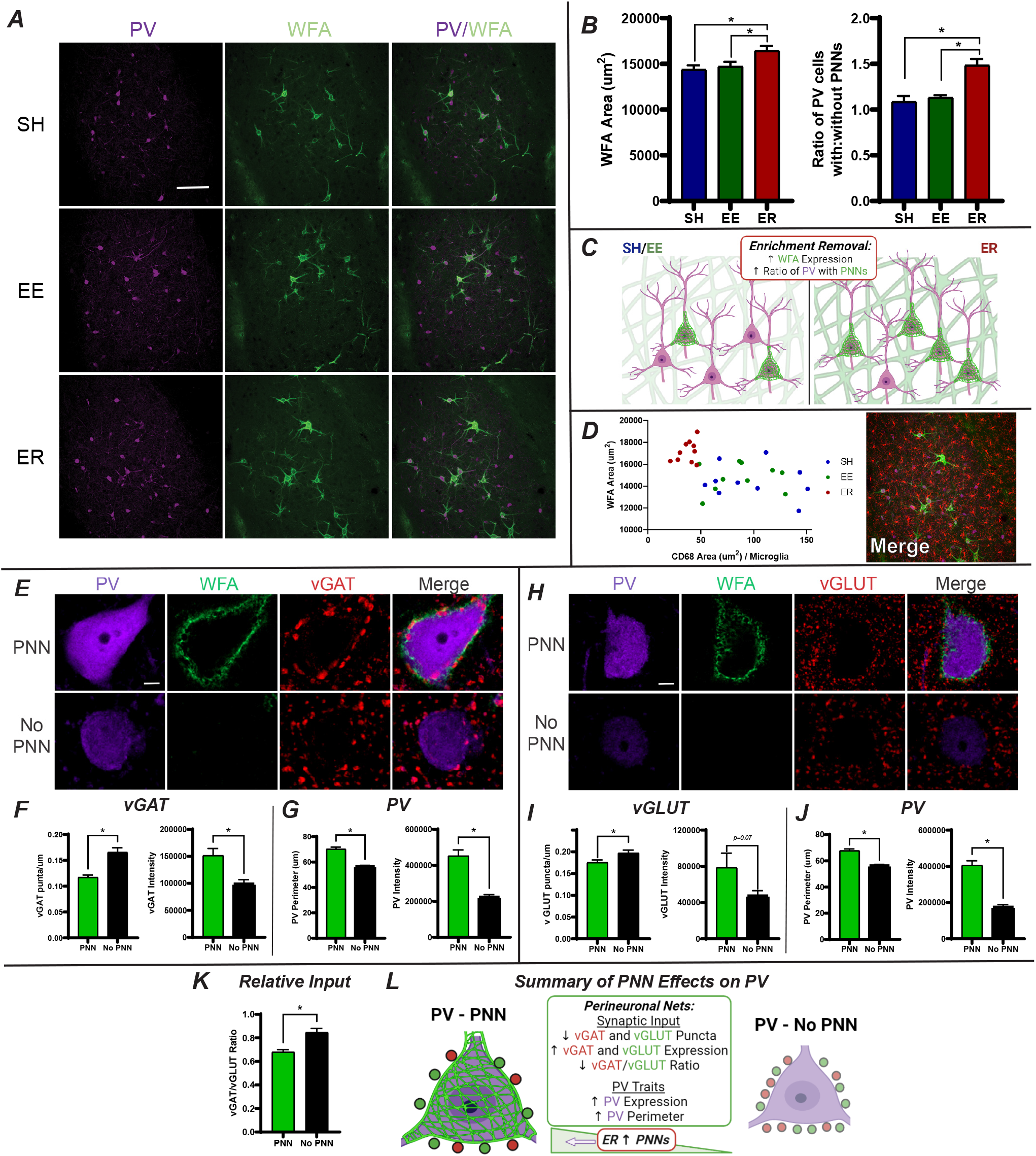
ER Increases BLA ECM and PNNs on PV Interneurons, Altering PV Synaptic Inputs and Phenotypic Characteristics. (A) Representative images of PV and WFA staining in the BLA (n=10/group; scale bar = 100 μm). (B) ER increases WFA area (μm^2^) and the ratio of PV cells decorated with PNNs. (C) Summary of ER effects on matrix phenotypes. (D) A significant correlation between WFA and CD68 suggests that microglia may be digesting the matrix less in ER, contributing to the present increases. Image shows IBA1/WFA/PV merge in the BLA. (E) Representative images of vGAT puncta surrounding PV cells with and without PNNs (n=10/group). (F) PNNs decrease vGAT puncta/μm and increase vGAT expression in those puncta. (G) PV cells with PNNs are larger and express more PV. (H) Representative images of vGLUT puncta surrounding PV cells with and without PNNs (n=10/group). (I) PNNs decrease vGLUT puncta/μm and increase vGLUT expression in those puncta. (J) PV cells with PNNs are larger and express more PV. (K) PNNs decrease the ratio of vGAT/vGLUT. (L) Summary of PNN effects on PV inputs and phenotypes. Given that ER increases PNNs, one would expect a shift in PV phenotypes in ER similar to those seen in PNNs here. * = p<0.05.

#### Connecting Microglia and the ECM

The omics suggested that the changes in ER microglia and ECM could be related. Microglia play a role in digesting the ECM, through mechanisms like integrin receptor signaling, matrix metalloprotease secretion, and phagocytosis^23,29,31,34^. Given that ER decreased microglia activity and increased ECM, it is possible that the loss of function in microglia could contribute to the buildup of ECM. We observed a significant correlation between microglial CD68 and WFA area. ER animals cluster together, demonstrating low CD68 area and high WFA area, supporting this potential link (Figure 4D).

#### PNNs Alter PV Synaptic Inputs and Characteristics

Increased BLA PNNs on inhibitory PV interneurons represents an avenue whereby the above changes can alter BLA output. PNNs regulate the physiology of PV cells^27,28,33^, and changes to PV could be indicative of altered BLA inhibitory tone^13,35^, an effect that could substantially impact ER behaviors. One of the major ways PNNs regulate PV is by altering synaptic input, acting as a physical guide for appositions onto PV^29,32^. Therefore, we tested whether ER-related increases in PNNs alter inhibitory (vGAT, Figure 4E) and excitatory (vGLUT, Figure 4H) synaptic inputs onto BLA PV cells. To eliminate sampling bias, we assessed vGAT and vGLUT terminals on equal numbers of PV cells with and without PNNs in each animal.

The presence of PNNs was associated with a decrease in both vGAT (Figure 4F) and vGLUT (Figure 4I) puncta on PV cells. However, puncta on PNN-bearing cells were significantly more intensely labeled, suggesting increased capacity for packaging of GABA and glutamate, respectively. Thus, these connections may be more efficient, favoring fewer, mature puncta over more numerous, but less developed puncta^36,37^. These effects were more robust in vGAT than vGLUT, a change made evident by the decreased vGAT/vGLUT ratio in PV cells with PNNs (Figure 4K). A decreased vGAT/vGLUT ratio indicates less relative inhibitory input onto these cells, suggesting that PNNs may allow for increased PV activity^38,39^. Differences in the PV cells themselves were observed, with PV cells with PNNs expressing more PV and having larger perimeters (Figure 4G,J). These changes are indicative of more mature PV cells with an increased calcium buffering capacity^40,41^, another sign of potentially more active PV cells with PNNs.

While ER did not affect these endpoints (Extended Data 7F,G), it is important to note that PNN-positive PV neurons were increased in ER relative to both control groups (Figure 4B). Thus, the impacts on synaptic efficacy and cellular maturity imparted by PNNs would disproportionately affect BLA PV neurons following ER (Figure 4L).

### Experiment 5: Expanding BLA Behaviors and Their Link to PNNs

We next explored the impact of ER on behaviors specifically linked to the BLA. We generated two new ER cohorts and conducted a series of behavioral tasks that are known for BLA involvement, including passive avoidance, social threat, acoustic startle, and cued fear conditioning^12,13,42–44^.

#### ER Impairs BLA-Related Behaviors

Two weeks after enrichment removal, rats received passive avoidance testing, followed by exposure to social threat in the three-chamber apparatus and subsequent assessment of acoustic startle (Figure 5A). No differences were detected between groups in the passive avoidance test^12,42^ (Extended Data 8A). In the social threat test, ER rats spent more time in the chamber where they had previously encountered a Long Evans retired breeder than EE controls (Figure 5B; Extended Data 8B), suggesting reduced fear memory relative to experienced animals (EE group) in a social context^43^. Finally, startle responses were enhanced in ER animals relative to both controls, consistent with prior data implicating the BLA in startle reactivity^13,44^ (Figure 5C; Extended Data 8C).

**Figure 5:**
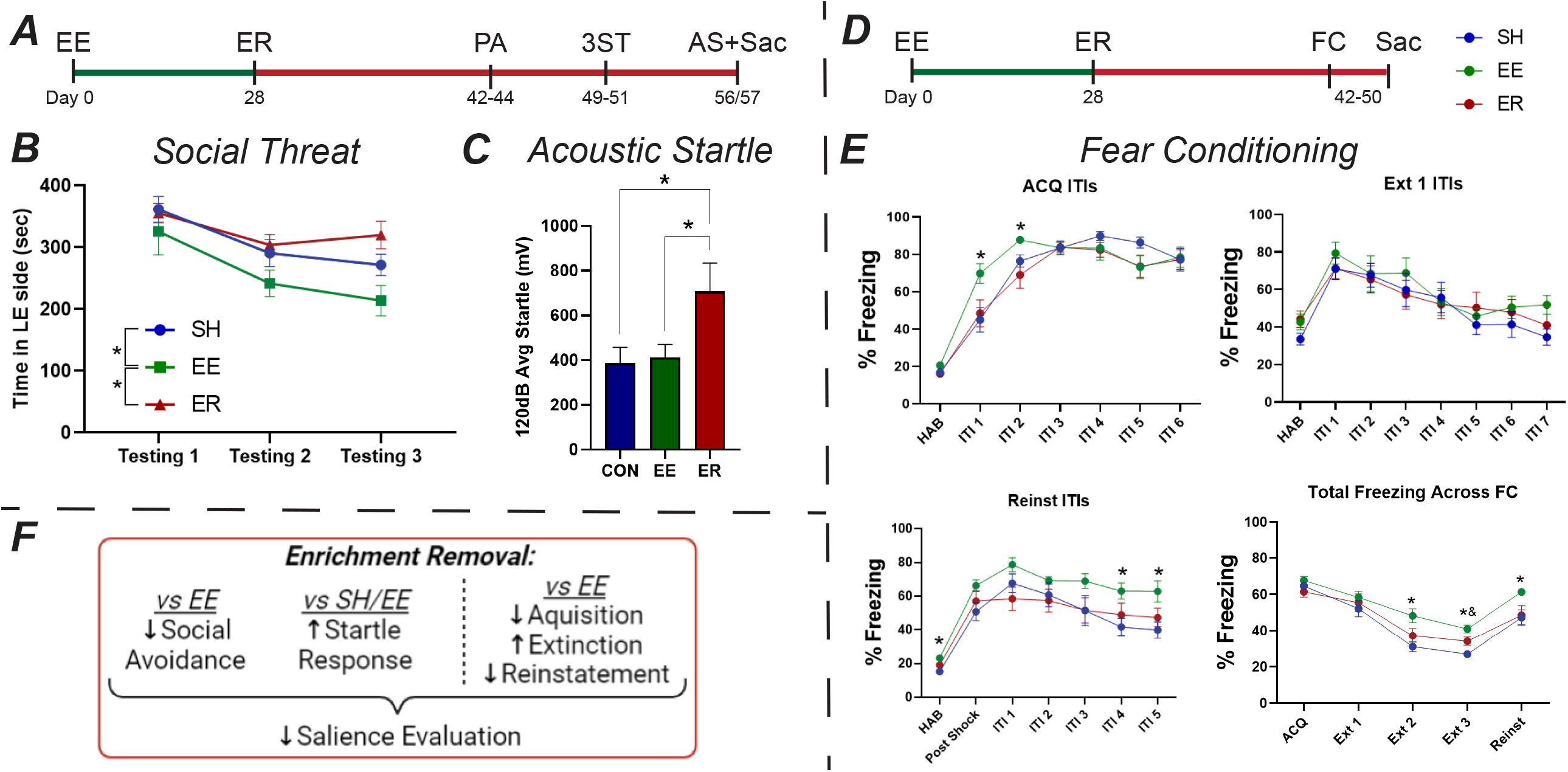
ER Impairs BLA-Related Salience Evaluation. (A) Timeline for passive avoidance (PA), three chamber social threat (3ST), and acoustic startle (AS) cohort (n=8-9/group). (B) Relative to EE rats, ER rats spend more time in the “threatening” side that previously housed an aggressive Long Evans rat. (C) ER rats show enhanced startle response. * = p<0.05. (D) Timeline for fear conditioning cohort (n=10/group). (E) Fear conditioning results. Relative to EE, ER rats exhibited delayed acquisition, faster extinction, and lower reinstatement. The first 3 panels show % freezing during ITIs on individual days, while the last shows total % freezing across all 5 days. * = p<0.05 for EE vs ER/SH. & = p<0.05 for ER vs SH. (F) Summary of behavioral results.

A second cohort was tested for cued fear conditioning and freezing responses (Figure 5D). ER rats acquired fear conditioning more slowly and extinguished more rapidly than EE controls, while also exhibiting reduced fear reinstatement (Figure 5E; Extended Data 8D). These results suggests that ER impairs fear learning and fear recall relative to EE in this more complex task^12,42^. There was no major difference evident between SH and ER groups, suggesting that ER actions are specific to loss of EE.

Taken together, these results suggest that ER generates a unique behavioral profile that is related to impaired fear recall and adaptability (Figure 6G). Both of these behavioral effects are consistent with a BLA that has attenuated plasticity and heightened inhibition^12,13^, tying these phenotypes back to the molecular effects observed above.

**Figure 6:**
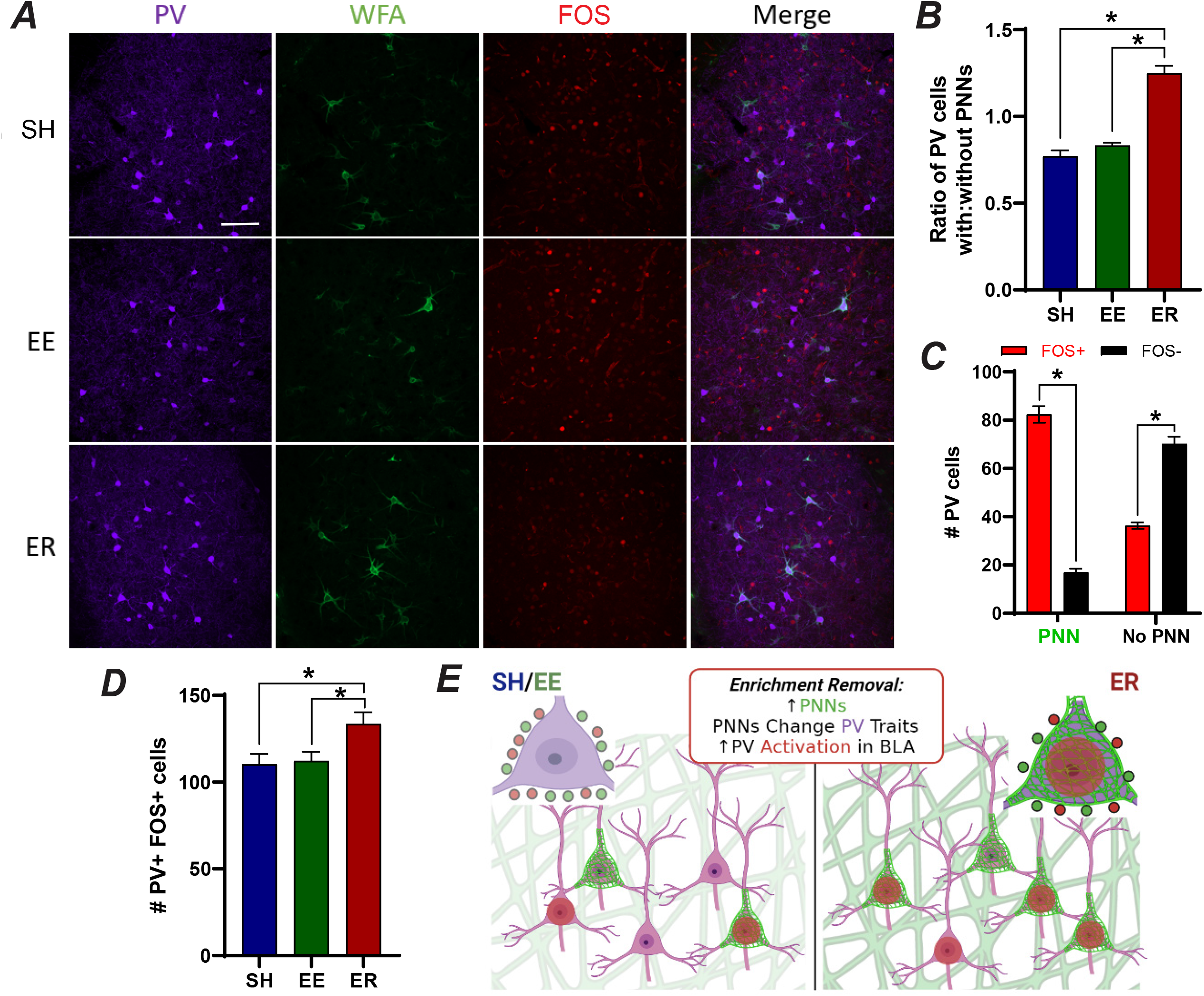
PV Interneurons are More Activated by Reinstatement in ER Rats. (A) Representative images of WFA/PV/FOS IHC (n=10/group). (B) Increased ratio of PV with PNNs in ER replicated in this cohort. (C) PV cells with PNNs were more likely to be FOS+ than PV cells without PNNs. (D) Together these differences resulted in more PV+ FOS+ cells in ER, indicating increased PV response to stress following ER. (E) Summary of FOS results and their connection to the above results regarding PNNs and PV. * = p<0.05.

#### ER Increases Fos Activation of BLA PV Cells Following Reinstatement

To test possible connections of ER-related behavioral changes to BLA activation, we assessed Fos induction following reinstatement testing (Figure 6A). The finding of increased PNNs on PV cells in ER rats replicated (Figure 6B), and expression of Fos was disproportionately enhanced in PV neurons surrounded by PNNs (Figure 6C). Importantly, Fos activation of PV neurons was increased in ER rats relative to both control groups (Figure 6D), further supporting the notion of increased PV-mediated inhibition in the BLA following ER (Figure 6E).

### Experiment 6: Testing the Necessity of BLA PNNs in Loss-like Behaviors

Lastly, we sought to establish a causal link between BLA PNNs and ER. Utilizing the enzyme Chondrotinase ABC (ChABC) which digests ECM and PNNs^23,33^, we investigated if blocking the increase in PNNs that occurs during ER could also block the development of loss-like behaviors.

EE and ER rats received bilateral injections of either ChABC or VEH into the BLA at the start of the removal period (Figure 7A) and received passive avoidance (PA), acoustic startle (AS), and forced swim (FST) testing. It is important to note that ChABC rapidly digests PNNs upon injection (∼90% depletion), and that PNNs then gradually repopulate over the next few months. In our hands, PNNs were still ∼60% depleted 3 weeks after injection (Figure 7B), so all behavioral testing was conducted within in this period.

**Figure 7:**
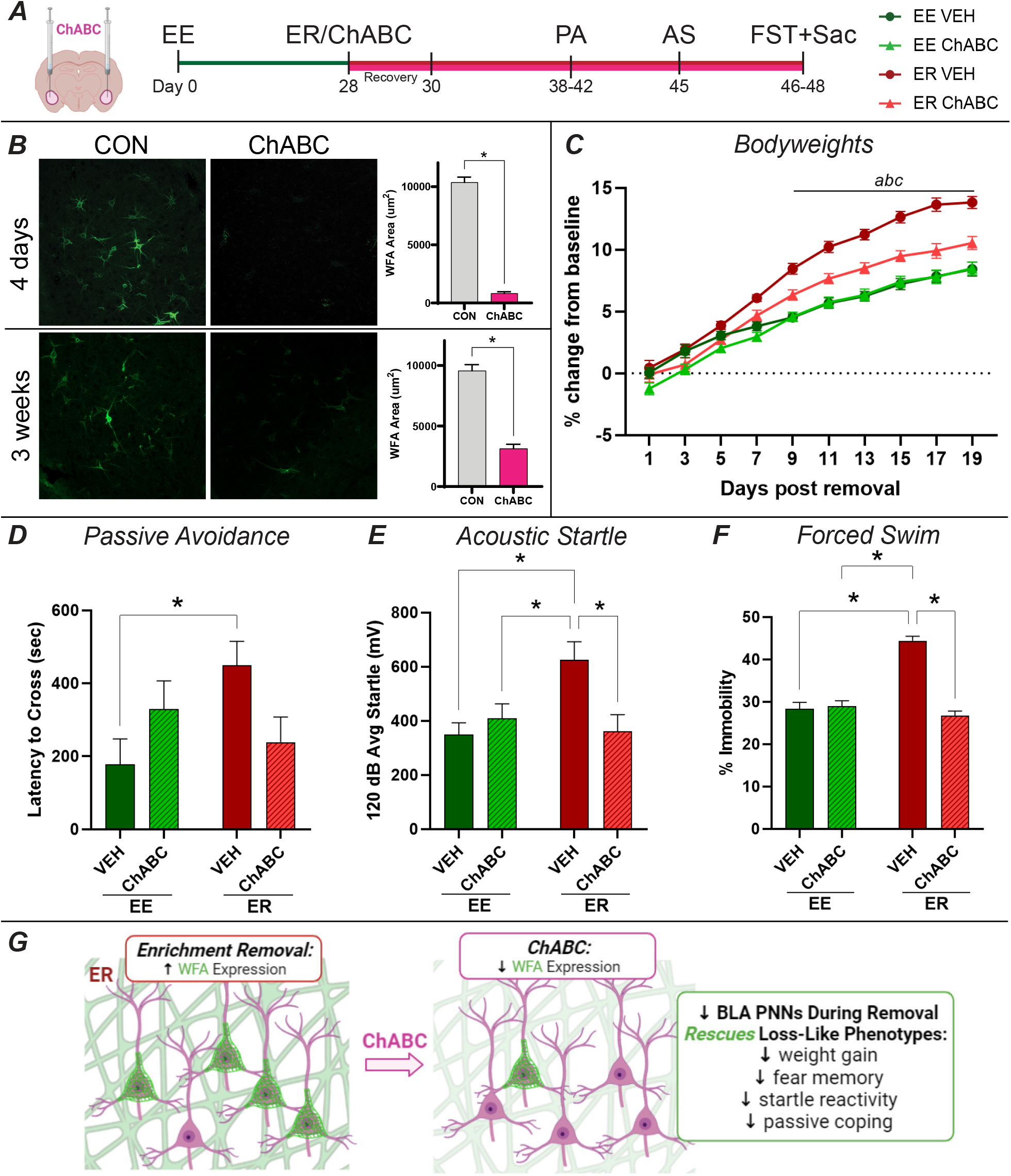
Depleting BLA PNNs Rescues ER Behavioral Phenotypes. (A) Timeline. Chondroitinase ABC (ChABC) was injected into the BLA at the time of removal to deplete PNNs. Rats were then subjected to passive avoidance (PA), acoustic startle (AS), and forced swim testing (FST). This study was a 2×2 design with housing (EE or ER) and treatment (ChABC or VEH) as factors. (B) Demonstration of ChABC effects on PNNs. PNNs are ∼90% depleted 4 days after ChABC injection and remain ∼60% depleted 3 weeks after injection. All behavioral testing was conducted within this timeframe to ensure that PNN levels would still be reduced to these levels. (C) Post-removal bodyweights show a blunting of ER effects in ER ChABC rats. While not to EE VEH levels, ER ChABC rats did gain significantly less weight than ER VEH rats. (D) ER VEH rats exhibited increased fear recall in passive avoidance testing relative to EE VEH rats, an effect that was blocked in ER ChABC rats. (E) Similarly, ER VEH rats showed a hyperresponsiveness to acoustic startle that was blocked by ChABC. (F) In the forced swim test, ER VEH rats exhibited increased immobility, while ER ChABC did not. (G) Summary of ChABC effects in ER rats. Overall, it appears that depleting PNNs in the BLA largely rescues loss-like phenotypes. * = p<0.05. a = p<0.05 for ER VEH vs EE VEH. b = p<0.05 for ER VEH vs ER ChABC. c = p<0.05 for EE VEH vs ER ChABC.

#### ChABC Rescues Multiple ER Phenotypes

One of the most established ER phenotypes is weight gain (Extended Data 1). ER rats that received BLA ChABC exhibited a blunted weight gain compared to ER VEH rats (Figure 7C). Behaviorally, ER VEH rats also exhibited multiple phenotypes that were blocked in ER ChABC rats. Latency to cross to the shock-conditioned side in PA was increased in ER VEH rats (Figure 7D; Extended Data 9A), the startle response was enhanced (Figure 7E; Extended Data 9B), and immobility was increased in the FST^45^ (Figure 7F; Extended Data 9C). In each case, ChABC was able to block the effects of ER, returning ER ChABC rats to the control levels.

Taken together, these results suggest that ChABC rescues multiple ER phenotypes (Figure 7G) and supports the necessity of BLA PNN accumulation in ER behavioral phenotypes.

## DISCUSSION

Psychological loss impacts most people at some point in their lives; however, the mechanisms underlying the experience of loss are poorly understood^1–3^. Here, we used enrichment removal (ER) in tandem with a series of comprehensive multi-omics, molecular, and behavioral approaches to investigate these mechanisms in the basolateral amygdala (BLA), as well as their subsequent functional consequences that contribute to loss-like phenotypes.

We first demonstrated that the BLA exhibited the strongest differential activation following ER. Then, multi-omics, spanning multiple cohorts, platforms, and analyses, indicated that BLA microglia and extracellular matrix (ECM) were dysregulated by ER. These relatively unexpected targets first emerged from hypothesis-free analyses, demonstrating the strength of our approach for identifying novel targets. The consistent implication of these targets across multiple levels affords us higher confidence that the changes are important to loss physiology. Further investigation supported these omics-derived hypotheses and revealed how ER dysregulates BLA microglia and ECM. ER decreased the size, complexity, and phagocytosis of BLA microglia, traits which suggest that ER microglia have an impaired capacity to surveille and interact with their surroundings (Figure 8A). While most studies assess conditions with overly active microglia and neuroinflammation, a loss of function in microglia can also be detrimental, as microglial monitoring and regulation of neurons and the microenvironment is critical to maintaining homeostasis. Indeed, depleting microglia can cause cognitive, social, and motor impairments^21,24–26^. Thus, the apparent decrease in microglia surveillance and phagocytosis following ER could attenuate the functional capacity of the BLA.

**Figure 8:**
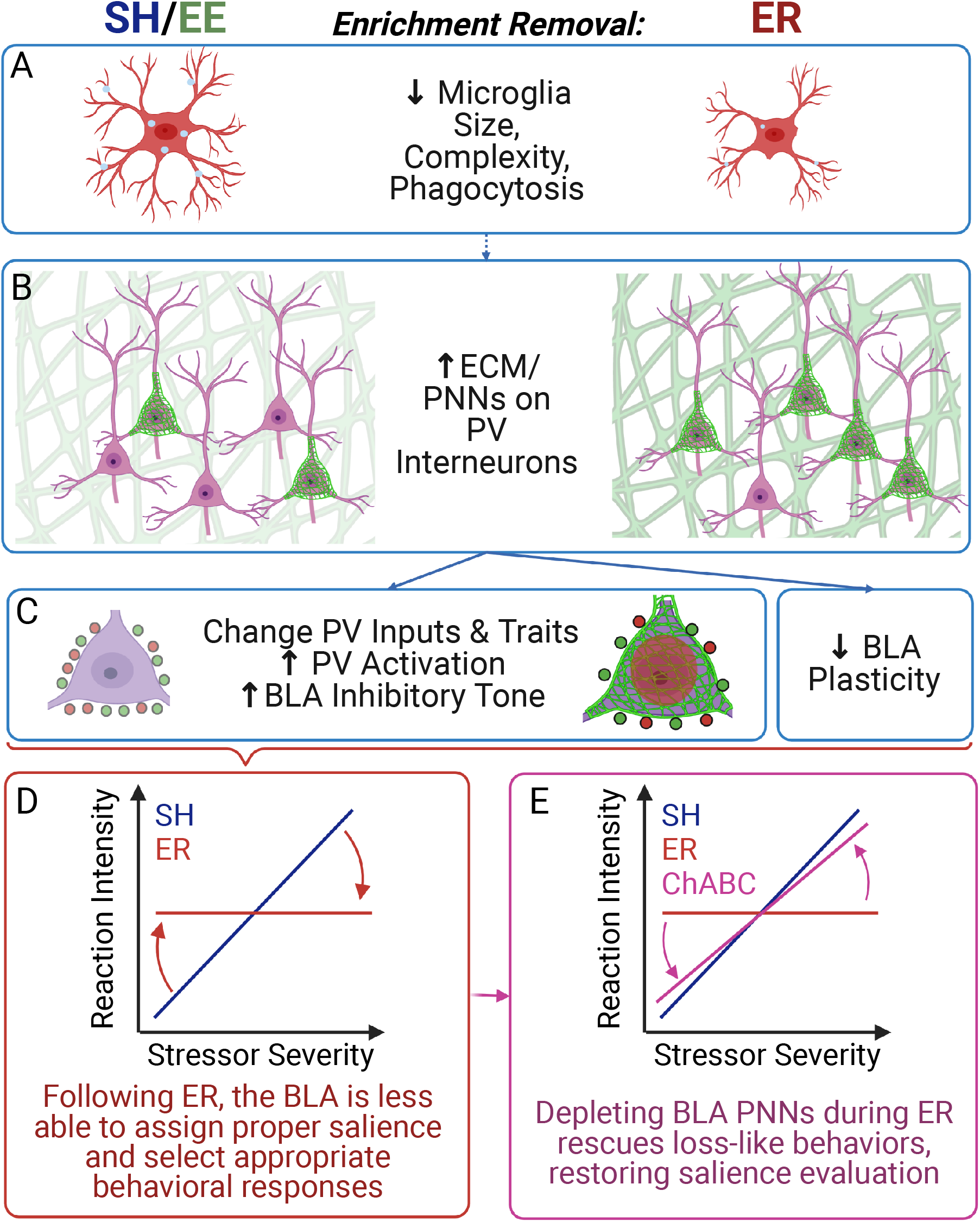
Summary of the Relationship Between BLA Microglia, ECM, and Loss-Like Behaviors. Collectively, the omics, molecular, and behavioral results presented here suggest the following: (A) ER leads to a loss of function in microglia that decreases their size, complexity, and phagocytic activity. (B) ER also increases ECM/PNNs in the BLA. These two findings may or may not be connected. (C) Increased PNNs influence PV interneurons in various ways, leading to increased PV activation and inhibitory tone in the BLA. Increased ECM in general is also expected to decreased plasticity within the BLA. (D) Together, these molecular changes appear to interfere with the BLA’s role in salience evaluation (triggering the proper type and level of response to various stimuli), uncoupling rats’ reactions from the intensity of the stressors and causing them to react too much or too little to various stimuli. (E) This effect can be largely rescued by depleting BLA PNNs. ER ChABC rats exhibit increased active coping (FST) and decreased fear responses (PA, AS) relative to ER VEH rats, suggesting that depleting PNNs at least partially restore the BLA’s salience evaluation capabilities. Ultimately, these complex mechanisms hold multiple similarities to loss in humans and represent several novel targets that could be used in developing therapeutics for people suffering from loss.

ER-associated changes in ECM expression and organization (increased PNNs decorating PV interneurons) are likely indicative of decreased plasticity, as PNNs form a physical barrier to new connections, making decorated cells less able to adapt (Figure 8B)^27,28^. Furthermore, we observed several PNN-associated changes to PV interneurons indicative of greater synaptic efficacy, less relative inhibition, and increased PV activity (Figure 8C). These findings align well the known roles of PNNs in protection from reactive oxygen species, participation in ion buffering, and organization of synaptic inputs^29,32^. Depleting PNNs is known to decrease PV activity and maturity^39,46^, supporting the conclusion that PV cells with PNNs are more active. Indeed, PV cells exhibited greater Fos responses to reinstatement in ER rats, suggesting that the BLA has greater inhibitory tone following ER. A less plastic, more inhibited BLA also supports the possibility of attenuated functional capacity, in line with the microglia findings. Finally, a correlation between microglial CD68 and WFA area suggests that these phenotypes may be linked. Microglia depletion can increase the amount of ECM^31,34^, so it is possible that the decreased microglia phagocytic marker seen here could lead to increased ECM deposition^23,31,34^ and a shift towards the PV phenotypes that accompany PNNs.

The functional consequences of these molecular changes in the BLA could contribute to loss-like behavioral phenotypes. Generally speaking, the BLA acts as an important neural node responsible for selection of appropriate behavioral responses to stimuli. It performs this role by gathering polysensory and associational information from numerous regions, gauging the salience and valence of those inputs, and signaling downstream regions responsible for appropriate responses. The BLA is key to evaluating both of these factors, and flexibility within the BLA is required for rapidly evaluating multiple inputs and switching responses over time^12,13,47–49^. Thus, decreased plasticity and increased inhibition following ER would be expected to interfere with its capacity to evaluate and select the appropriate response to various stimuli. ER rats showed exaggerated responses to some stimuli (startle) while showing blunted responses to others (fear conditioning), suggesting impaired evaluation of salience (Figure 8D)^12,13,48^. For example, startle responses may be enhanced by BLA inhibition, losing the ability to appropriately gate response to the relatively low salience threat. In contrast, responses to high salience threats requiring freezing or social avoidance are attenuated. Our data align with previous studies noting that depletion of BLA ECM increased behavioral flexibility and adaptability^23,33^. We functionally tested this relationship by depleting BLA PNNs during the removal period. This manipulation successfully rescued multiple ER physiological and behavioral phenotypes, blunting weight gain and blocking enhanced avoidance, startle, and passive coping. These findings support the necessity of BLA PNN accumulation in driving behavior following ER, suggesting an intriguing target for ameliorating loss. This suggests that blocking the increase in PNNs that occurs during ER can, at least partly, restore an animal’s capacity for salience evaluation, possibly via increased plasticity and dampened inhibition. Interestingly, ChABC had minimal effects in EE rats, suggesting that there is a unique interaction between the state of loss and PNN depletion.

It is important to note that the ER group diverges from both EE and SH groups on most endpoints, indicating that the loss of EE is not a simple return to a ‘basal state’ but is a unique condition generated by EE loss. Understanding how loss affects females was not addressed in these studies, due in part to a different behavioral and omics signatures observed in ER females and will need to be a major emphasis moving forward. Future studies will be required to determine how changes in PV activity in PNN-bearing cells affect output and oscillatory activity within the BLA, as well as to further evaluate the relationship between changes in microglial function and ECM hypertrophy.

Humans suffering from loss often exhibit emotional blunting and impaired adaptability^1,2,6^. This emotional rigidity aligns well with the mechanisms proposed here, namely that loss of a motivationally significant stimulus impairs an individual’s salience evaluation, resulting in a mismatch between stimuli and response. These experiments put forth BLA PNNs an intriguing target for ameliorating loss symptoms in humans. In addition, symptoms of atypical depression (often precipitated by loss) are associated with decreased BLA activity and responsiveness^1,50^, suggesting that the local loss-related neurobiological changes in microglia, ECM, and PV interneurons observed here could have broader implications for a range of mood disorders. Further understanding the neurobiology of loss will be essential for development of both behavioral and pharmaceutical intervention approaches to limit its nearly universal impact on health and well-being.

## METHODS

### Subjects

Adult male Sprague-Dawley rats were used for all experiments. Rats were obtained from Envigo (Indianapolis IN USA) at 8 weeks of age and given 1 week to acclimate to the vivarium prior to experimental manipulations. The vivarium was temperature and humidity controlled, and chow and water were provided ad libitum. There was a 12-hour light cycle (lights on at 8:00 AM, lights off at 8:00 PM). All procedures were conducted in compliance with the National Institutes of Health Guidelines for the Care and Use of Animals and approved by the University of Cincinnati Institutional Animal Care and Use Committee.

### Enrichment Removal Protocol

Rats were randomly assigned to 3 housing conditions: standard housing (SH), environmental enrichment (EE), and enrichment removal (ER). SH rats were kept in standard housing conditions, which was either single or pair housing as noted below. EE rats remained in enrichment housing throughout the entire protocol. Enrichment housing consisted of 10 same-sex rats housed in a 1m^3^ multi-level wire mesh cage with 5-6 toys and shelters that were rotated weekly (Figure 1A). Toys included plastic huts, tunnels, balls, chains, and objects such as small shovels, pitchers, and buckets. While rats obtained physical enrichment from climbing and play, enrichment housing does not contain a running wheel. As in previous studies^8,9^, EE rats were kept in this enrichment housing during their active cycle (8 PM to 8 AM), and then moved to single housing during the day (8 AM to 8 PM) to permit food and water intake measurements and behavioral analyses. ER rats were maintained in the same conditions as EE rats for 4 weeks. ER rats were then removed from EE and placed into 24-hour single housing for the remainder of the experiment, with no further enrichment. SH rats were handled at the same times as EE and ER rats. For molecular experiments, rats were kept naïve to behavioral testing and maintained on ER for 2 weeks prior to sacrifice (Figure 1A). This ensures that ER phenotypes have sufficient time to incubate, as was suggested in our prior studies^8^.

### Experimental Design

Six experiments were conducted for the present analyses (Table S1). All experiments utilized the ER protocol, and procedural differences between studies will be noted where appropriate. Note that in all experiments ER rats gained more weight than EE and SH rats, demonstrating the consistency of the model (Extended Data 1; Figure 7C). The experiments were conducted as follows:

### Experiment 1: Fos Expression Screen

Fos expression profiling was first used to determine stress-responsive brain regions that showed differential activation in response to EE and ER exposure (n=10/group) to act as a guide for the upcoming omics studies. In this case, SH rats were kept in single housing (previous experiments did not document significant differences between single housing or pair housing on physiological or behavioral endpoints in male rats^8^), and EE rats were kept in enrichment housing 24 hours per day. These animals also experienced behavioral testing in the form of an open field test, social interaction test, and forced swim test (FST) starting 1 week after removal, conducted in 1-week intervals. Animals were sacrificed and perfused 120 minutes after the FST to assess Fos activation to acute stress.

#### Immunohistochemistry

Rats were injected with an overdose of sodium pentobarbital (150 mg/kg) 120 minutes after the initiation of the FST and then transcardially perfused with 0.9% saline followed by 4% paraformaldehyde in phosphate-buffered saline (PBS) solution. Brains were removed and placed in 4% paraformaldehyde in PBS overnight, after which they were placed in 30% sucrose solution in phosphate-buffered saline (4 ºC) until sectioning. Brains were cut into 35μm serial sections using a sliding microtome and stored at -20ºC in cryoprotectant solution (0.1 M phosphate buffer, 30% sucrose, 1% polyvinylpyrrolidone, and 30% ethylene glycol). Sections were removed from cryoprotectant, rinsed 5 × 5 min with 50mM KPS, treated with 1% hydrogen peroxide for 10 min, rinsed 5 × 5 min with KPBS, and treated with 1% sodium borohydride (Fisher Scientific). After incubation in sodium borohydride, sections were rinsed 5 × 5 min with KPBS, and incubated for 1 hour in blocking solution (50mM KPBS, 0.1% bovine serum albumin, 0.2% Triton X-100). From blocking solution, sections were transferred into wells with primary mouse anti-cFOS antibody (Santa Cruz, no. sc-52) (1:5000 in blocking solution) for incubation overnight. On the second day sections were rinsed 5 × 5 min with KPBS, incubated for 1 hour in biotinylated anti-mouse secondary antibody (Vector Laboratories Inc.), and rinsed 5 × 5 min in KPBS. Following secondary antibody incubation, sections were incubated for 1 hour in avidin-biotin complex (Vector Laboratories Inc.) (1:800 in 50mMKPS + 0.1% bovine serum albumin), rinsed 5 × 5 min in KPBS, and incubated for 10 min in 0.02% 3, 3-diaminobenzidine (Sigma Aldrich) with 0.05% hydrogen peroxide. To terminate DAB incubation, sections were rinsed 4 × 5 min with KPBS. Finally, sections were mounted in 50mM phosphate buffer, dehydrated with an ethanol series (0%, 50%, 75%, 95%, 100%), incubated in xylene for a minimum of 7 minutes and coverslipped with DPX (Sigma Aldrich).

#### Image Analysis and Fos Quantification

Images were obtained using Zeiss Imager Z.1 (Carl Zeiss Microimaging) and 10x objective. The BLA, medial amygdala (MEA), central amygdala (CEA), infralimbic prefrontal cortex (IL), prelimbic prefrontal cortex (PL), bed nucleus of the stria terminalis (BST), paraventricular nucleus (PVN), nucleus accumbens (NAc), and dentate gyrus (DG) were identified using a rat brain atlas^14^. When possible, 3 images were taken from each brain region (rostral, mid, caudal). Images were quantified using Scion Image. A consistent threshold was used for all images to select and count immunolabeled cells in each ROI. Cells were counted bilaterally and averaged across sections. The total number of Fos positive cells are expressed per arbitrary unit area.

#### Statistics

Group effects were analyzed by one-way ANOVA using Sigma Stat. All post hoc testing utilized Fisher’s Least Significant Difference (LSD). Outliers were determined by values that fall outside the mean ± 1.96 times the standard deviation.

### Experiment 2: Case-Control RNAseq

Based upon the results of the Fos screen, the BLA was selected for further analysis with RNAseq, a method which provides a comprehensive picture of the transcriptional landscape of a tissue. This experiment was conducted on a new cohort of male rats (n=10 per group), and the ER protocol was run with active cycle enrichment as described above. SH rats were kept in single housing. SH, EE, and ER rats were sacrificed by rapid decapitation 2 weeks after removal (6 weeks of active cycle enrichment for EE group). Brains were collected, flash frozen in isopentane, and stored at -80□C.

#### Micropunch Collection of the BLA

Brains were blocked at the base of the cerebrum, mounted on the cryostat, and cut in 500 μm sections. Sections were mounted on slides and 1mm diameter micropunches were collected bilaterally from 3 sections containing BLA (Figure 1C; AP -1.5 to AP-3.0)^14^. Micropunches were stored at -80□C. Six rats per group were used for further analysis based on cost and punch placement.

#### RNA Extraction

For each sample, RNA was obtained using Qiagen RNAqueous-Micro isolation kits. RNA quantity and quality were assessed using a Nanodrop ND-1000 spectrophotometer.

#### RNAseq Protocol

RNAseq was performed by Genomics, Epigenomics and Sequencing Core (GESC) at the University of Cincinnati. Briefly, ∼400ng RNA per sample underwent polyA RNA isolation, library preparation, and cluster generation prior to sequencing on the Illumina HiSeq system. The sequencing conditions were single end 50 bases and 25 M reads per sample. Bioinformatics analyses of this data is detailed below.

### Experiment 3: Validation and Expansion

In order to gain a more complete understanding of the molecular mechanisms suggested by Experiment 2, these studies were expanded to other omics platforms. In this case, RNAseq, shotgun proteomics, and serine-threonine kinomics were run in parallel on the same tissue sample. These parallel analyses were run on pooled tissue samples from the BLA. This triple-omics approach was used to examine protein expression and kinase activity in addition to RNA expression. This approach not only served as a validation of Experiment 2 (to determine reproducibility between cohorts), but also allowed us to examine ER-induced changes at multiple levels, providing more complete molecular signatures. In this case, SH rats were kept in pair housing, as it serves as a more realistic control condition and enables future studies of sex differences^9,8^. The ER protocol was conducted as describe above (n=10 per group), and rats were sacrificed by rapid decapitation 2 weeks after ER. Micropunches of the BLA were collected as described in Experiment 2.

#### Tissue Processing: The Triple Prep

In order to examine RNA expression, protein expression, and kinase activity in the same tissue sample, we developed the “triple prep” protocol (Figure 1C). As opposed to splitting a tissue sample for different analyses, this “triple prep” involves homogenization of the sample and extraction of RNA and protein simultaneously. This step should guard against observed differences between analysis platforms stemming from regional heterogeneity within a sample.

Micropunches were homogenized in 50 μl of MPER, Halt, and RNAse inhibitor (ThermoFisher) using a hand pestle. This homogenized sample was then split into 3 portions for RNA and protein extraction. Twenty-five μl of homogenate was used for RNA extraction, which was performed using Qiagen Mini Kit according to the manufacturer’s protocol. Amount and quality of RNA was determined using a Nanodrop. All 260/280 values were greater than 1.8. Twenty-five μl of homogenate was maintained for protein samples, and subsequently split for proteomics and kinomics analysis. The proteomics samples were used directly, while the kinomics samples were subjected to 10 minutes of centrifugation at 14,000 rpm and 4ºC. Supernatant was collected for the kinomics samples. The amount of protein present in both proteomics and kinomics samples were assessed via BCA assay. All 3 types of samples were stored at -80ºC.

Once all extractions were complete, pools were made for each sample type and experimental group (n=10 individuals per pool). The logistics of using 3 platforms prohibited us from running these as individual samples, so representative pool samples were generated for each group for each analysis. For example, this generated samples such as “Male-ER-BLA” that were comprised of combined samples from 10 rats. “Amount” of RNA/protein was comprised of equal representation by each individual sample. Once pools were generated, RNA samples were submitted to the CCHMC DNA Sequencing Core for RNAseq, protein samples were submitted to the UC Proteomics Core for LCMS shotgun proteomics, and protein samples were submitted to the Toledo Core for serine threonine kinomics.

#### RNAseq Protocol

Quality of RNA samples was assessed using an AATI Fragment Analyzer. All samples had an RQN greater than 8. Samples then underwent RNA polyA stranded library preparation. RNA sequencing was then performed via HiSeq 2500 Rapid Sequencing. The sequencing conditions were paired end 75 bases and 20 M reads per sample.

#### Proteomics Protocol

Liquid Chromatography Mass Spectrometry (LCMS) was used to run data-dependent acquisition (DDA) shotgun proteomics on the pooled protein samples. Three technical replicates were run for each sample (SH, EE, ER); however, due to technical difficulties only one replicate per group yielded usable data that underwent subsequent analyses.

#### Kinomics Protocol

The pooled protein samples were run in triplicate on PamGene Serine Threonine Kinase (STK) Arrays. This microarray approach measures phosphorylation of 144 peptides, and then bioinformatics is used to infer upstream kinase activity. One technical replicate of each sample (SH, EE, ER) was run on each STK chip, and 3 chips were run in total.

### Bioinformatics Analyses of Omics Data from Experiments 2 and 3

Experiments 2 and 3 both yielded vast amounts of data regarding the molecular signatures of the BLA following EE and ER. Our primary goal here was to identify targets and pathways that replicated across cohorts and omics platforms, with replication lending greater assurance that those target/pathways play important roles in ER. For example, we can have greater confidence that a phenotype that is seen at the RNA, protein, and activity levels (and replicates across multiple RNA cohorts) is contributing to ER. As such, we used various bioinformatics tools to summarize the data into meaningful functional profiles that could be easily compared (Extended Data 3). We also developed several novel tools to aid in this effort that are discussed below. Altogether, these analyses allow us to better define and compare molecular landscapes of the BLA to identify key targets and pathways that could be driving ER and potentially targeted by novel therapeutic strategies to ameliorate loss.

### RNAseq Analysis

Bioinformatic analyses of RNAseq data were conducted in the same manner for Experiments 2 and 3. Raw fastq files were obtained from respective cores. Reads were aligned using HISAT2 and normalized using DESeq2. Fold change values were then calculated for all detected protein-coding genes with an average base mean above 1. These differential expression values were generated for 3 contrasts (EEvSH, ERvSH, ERvEE) which were then pushed through our comprehensive analysis pipeline to allow for confirmation across analysis platforms. This pipeline functions on the same principle driving the multi-omics approach: targets/pathways obtained consistently across different analyses are more likely to play a true role in the present phenotype. This method also guards again biases in single platforms that could mask important effects or point to false leads. Our pipeline (Full Transcriptome Pathway Analysis, Targeted Pathway Analysis, and Signature Analysis) is detailed below.

It is important to note that the following analyses are based on fold change (FC), rather than statistical significance. Given that Experiment 3 was run on pooled samples, we were unable to perform statistics on these datasets and needed to rely on fold change to assess differential expression. Experiment 2 was also analyzed in this way to maintain consistency between experimental analyses. This technique also emphasizes the functional implications of observed changes, which aligns with the goals of this project to identify biological substrates of loss.

#### Full Transcriptome Pathway Analysis

The first analysis utilized Gene Set Enrichment Analysis^15^ (GSEA; gsea-msigdb.org) to assess transcriptome-wide pathway enrichment. This technique considers all data and detects subtle changes impacting the full transcriptome. GSEA uses a rank-ordered list of all detected transcripts (ordered by fold change (FC)) and compares that order to established pathway gene sets. If the genes in a pathway are mainly high on the FC list, that pathway is positively enriched. Negatively enriched pathways show a high concentration of genes at the bottom of the FC list. GSEA was conducted using the fgsea R package. RNAseq data was sorted by FC rank order and compared to GO pathways obtained from BaderLab (download.baderlab.org/EM_Genesets/). Gene sets used in this analysis contained between 15 and 500 genes, and GSEA was run with 10,000 permutations. GSEA generates p values and enrichment scores (ES) to assess the significance and strength of this enrichment. Pathways with p<0.05 were extracted and condensed using a new tool we developed called PathwayHunter, which uses semantic similarity to sort pathways into functionally similar categories^16^.

While GSEA is a great tool for revealing subtle transcriptome-wide changes, it yields thousands of pathways, making it difficult to ascertain functional meaning from the changes. PathwayHunter was designed to take those thousands of pathways and summarize them into enriched categories. These categories were initially generated from comparison of the semantic similarity of the 44,000 Gene Ontology pathway titles and descriptions, which sorted them into ∼500 clusters. PathwayHunter then uses hypergeometric overlap to determine which of these clusters are overrepresented in the significant pathways from the dataset. It assigns a -log10 p value to demonstrate the strength of this overrepresentation (with higher scores indicating stronger involvement), along with a 2-word label generated from the semantic similarities to describe the category. While this tool is a great first step in narrowing down the thousands of significant pathways obtained from GSEA, an additional step of manual curation based on *a prior* knowledge was used to more broadly group together the overrepresented categories into functional themes. Themes were based on the PathwayHunter label as well as the pathway titles/descriptions contained within that category. These themes can then be examined with the GSEA ESs to determine which types of pathways were altered in which direction across the experimental groups. This process of condensing and categorizing enriched pathways enabled us to gain a broader understanding of the molecular changes associated with EE and ER.

Additionally, leading-edge genes, which are genes that are driving the observed pathways, were collected from GSEA. Cell-type enrichment of top 100 upregulated and downregulated leading-edge genes was then conducted in Kaleidoscope, using the “Brain RNA-seq module,” to determine if certain cell types were preferentially impacted by ER.

#### Targeted Pathway Analysis

The second analysis utilized Enrichr^17^ (maayanlab.cloud/Enrichr/) to assess pathway enrichment within the top 100 differentially expressed genes (DEGs) by FC. This is a more traditional pathway analysis and is more targeted to detect pathways with strong enrichments. The top 100 upregulated DEGs were extracted and entered into the Enrichr R package, where significantly (p<0.05) upregulated pathways were collected for GO Biological Process, GO Molecular Function, and GO Cellular Component. The same analysis was done for the top 100 downregulated genes, and GO terms were condensed by PathwayHunter as described above. By condensing pathways from both full and targeted pathway analyses with PathwayHunter, we were able to make direct comparisons between the two to determine which functional categories were implicated by both analyses.

#### Signature Analysis

The third analysis utilized iLINCS^18^ (ilincs.org/ilincs/), which is a bank of L1000 signatures for various cell line manipulations, to identify perturbagens related to ER signatures. The L1000 is a set of 978 genes, whose expression is known for thousands of cell line signatures upon exposure to various perturbagens. The L1000 was extracted from our RNAseq data and uploaded to iLINCS for comparison to this data bank. The top 20 discordant perturbagens were determined by concordance score. These discordant perturbagens induce a signature which is opposite of that induced by ER, thus the drug in theory could reverse some of the molecular effects of loss. The top 20 concordant perturbagens, whose signatures theoretically would recapitulate ER, were also collected to inform on pathways involved in ER. Mechanism of action (MOA) was determined for these perturbagens using the L1000 Fireworks Display. These top perturbagens were then compared with the full and targeted pathway results to determine which consistent pathways were further implicated by MOA.

### Proteomics Analysis

Differential expression values were calculated for peptides that were detected with 99% confidence in all groups and subjected to quartile normalization. These peptides were converted into their corresponding proteins using PiNET (eh3.uc.edu/pinet/peptideToProtein). The top 100 upregulated and downregulated proteins (by FC) were then input into Enrichr for pathway analysis. Enrichr was chosen to allow for easier comparison with the RNAseq and because the signatures were not large enough to benefit from GSEA. Significant pathways were condensed using PathwayHunter and results were compared to the corresponding RNAseq results.

### Kinomics Analysis

Phosphorylation levels were obtained for the 144 peptides on the STK chip. Kinome Random Sampling Analyzer^20^ (KRSA) was used to infer kinase activity from this data. This approach uses a permutation method to generate z-scores for each kinase. In order to correct for between-chip differences, KRSA was run separately for each chip and the resulting z-scores were averaged for each kinase. Kinases with a z-score > 1.5 were considered to be over-represented, and kinases with a z-score <-1.5 were considered to be under-represented. Additionally, individual peptide phosphorylation was examined for targets of interest that emerged from the RNAseq analysis. Pathway analysis was run using Enrichr on the top differentially phosphorylated peptides (FC > 1.2) and condensed using PathwayHunter to enable functional comparison between the omics platforms.

### Cross-Platform Summary

As described above, the primary goal of these multi-level omics analyses was to identify commonly implicated targets and pathways (Extended Data 3). Direct comparison between the RNAseq (pathway analyses from Experiments 2 and 3), proteomics, and kinomics was enabled by PathwayHunter, which revealed functional categories that were implicated across platforms, analyses, and cohorts. These categories were further defined and bolstered by the perturbagen, cell-type, and upstream kinase results, which pointed to more specific targets within these themes. These commonly implicated results formed the basis for novel hypotheses that were investigated in subsequent cohorts.

### Experiment 4: Hypothesis Investigation: Microglia and the Extracellular Matrix

In order to validate hypotheses derived from the omics experiments, we generated a new cohort of ER rats (n=10/group) that were sacrificed and perfused 2 weeks after removal, as described above (Experiment 1). Brains were cut into 40 μm serial sections and stored at -20ºC in cryoprotectant to allow for immunohistochemical (IHC) analysis of implicated endpoints.

#### Antibodies

Further investigation was conducted on microglia and the extracellular matrix (ECM). IBA1 (Wako, rabbit anti-IBA1, 019-19741, 1:1000), a common microglial marker, was used to determine microglia counts and morphology. CD68 (Abcam, mouse anti-CD68, ab31630, 1:500) was used to determine the level of microglia phagocytosis. Wisteria Floribunda Agglutinin (WFA) (Vector, biotinylated, B-1355, 1:1000) was used to label chondroitin sulfate proteoglycans (CSPGs), the primary component of the ECM. Given that a major element of the ECM is perineuronal nets (PNNs), which primarily surround Parvalbumin (PV) interneurons, we also stained for PV (Sigma, mouse anti-PV, P3088, 1:2000). Follow-up studies utilized vGAT (Synaptic Systems, rabbit anti-vGAT, 131 002, 1:1000) and vGLUT (Synaptic Systems, rabbit anti-vGLUT, 135 303, 1:2000) to examine inhibitory and excitatory synaptic inputs onto PV cells, respectively. Secondary antibodies included Cy3 goat anti-rabbit (Thermo A10520), Streptavidin AlexaFluor 488 (Thermo S11223), and Cy5 goat anti-mouse (Thermo A10524) and were all run at 1:500.

#### Immunohistochemistry

Four triple label IHCs were run: IBA1/WFA/PV, IBA1/WFA/CD68, WFA/PV/vGAT, and WFA/PV/vGLUT. IHCs were run using the following general protocol and above antibody concentrations. Sections were removed from cryoprotectant, rinsed 5 × 5 min with 1xPBS, blocked for 1 hour (0.1% BSA and 0.4% Triton in 1xPBS) at room temperature, then incubated in primary antibody in block overnight at 4ºC. Sections were then rinsed 5 × 5 min with 1xPBS before being moved into secondary antibody in block for another overnight incubation at 4ºC. Sections were again rinsed at 5 × 5 min with 1xPBS before mounting. Slides were allowed to dry overnight before coverslipping with gelvatol (Sigma Aldrich, 10981). The only variation of the protocol was increasing the block for the CD68 run (blocking: 1% BSA, 5% NGS, and 0.4% Triton; antibody block: 1% BSA, 2.5% NGS, and 0.4% Triton).

#### Image Analysis and Quantification

Images were obtained using a Nikon C1 confocal microscope (Nikon Instruments Inc). For initial microglia and ECM analyses, z-stacks (0.4 μm steps) of the IBA1/WFA/PV IHC were captured using the 20x objective. Images were taken bilaterally from 3 levels of the BLA (approx. at AP -2.0, -2.5, and -3.0), flattened into maximum intensity projections, and analyzed using Image J. Image settings were kept constant for all animals. All counts and measures were collected by a blinded observer. IBA1 endpoints included number, area (μm^2^), and circumference (μm) of microglia. WFA endpoints included WFA area (μm^2^), WFA intensity, and the ratio of PV cells with and without PNNs. These counts determined how many PV cells were decorated with PNNs, and additionally counted total PV cells and PNNs in the BLA.

For microglia morphology and phagocytosis analyses, z-stacks (0.2 μm steps) of the IBA1/WFA/CD68 IHC were captured using the 40x objective with a 2x zoom and included 2 bilateral levels of the BLA (approx. at AP -2.0, -2.5). CD68 area (μm^2^) and area per microglia were quantified using ImageJ. Microglia morphology was assessed using the filament tracer tool in Imaris (n=8 cells/animal), with primary endpoints including process area, process volume, number of branches, and number of terminal points.

For synaptic inputs analyses, z-stacks (0.2 μm steps) of the WFA/PV/vGAT and WFA/PV/vGLUT IHCs were captured using the 60x objective and included 2 bilateral levels of the BLA (approx. at AP -2.0, -2.5). vGAT and vGLUT puncta apposed to PV cells were counted manually by a blinded observer using ImageJ and the Hyperstacks plugin. Given that this analysis was designed to determine the impact of PNNs on PV inputs, equal numbers of PV cells with and without PNNs were counted (n=8 PV cells/animal, 4 with PNNs, 4 without PNNs). Puncta within 1 μm of the cell’s perimeter were considered to be apposed, and puncta were counted at 3 levels for each cell^41^. Counts are presented as puncta/μm of PV perimeter. Additional endpoints measured for both vGAT and vGLUT included puncta intensity, PV intensity, and PV perimeter. vGAT/vGLUT ratio was also calculated.

#### Statistics

Group effects were analyzed by one-way ANOVA using GraphPad Prism 9. All post hoc testing utilized Tukey tests. Outliers were determined by values that fall outside the mean ± 1.96 times the standard deviation. Pearson correlations were used to explore relationships between microglia and ECM endpoints.

### Experiment 5: Expanding BLA-Related Behavioral Profiling Following ER

The data thus far have indicated that ER induces multiple molecular changes in the BLA. However, the behaviors that we have previously characterized in ER are not very specific to the BLA. In order to tie our molecular results to behavioral outcomes, we generated two new cohorts of ER rats to 1) expand our behavioral profile of ER to behaviors known to centrally involve the BLA and 2) examine activation of BLA PV cells following behavioral testing.

### Behavioral Testing and Analysis

Two cohorts of ER rats were generated the ER protocol as described above (Figure 1A). Behavioral testing began 2 weeks after removal. In the first cohort, testing consisted of passive avoidance^12,42^, 3 chamber social threat^43^, and acoustic startle^13,44^ (Figure 5A). In the second, testing consisted of cued fear conditioning^12,42^ (Figure 5D). The BLA is known to play a central role in the behaviors tested. All behaviors were conducted between 8:00 and 14:00. All rats were sacrificed 90 minutes after the final behavioral task.

#### Passive Avoidance

Passive avoidance was run using the Gemini system (San Diego Instruments) and consisted of 5 phases: habituation, training, and 3 days of testing. The passive avoidance chamber features a dark side and a light side divided by a door. The door was open for the habituation phase (day 1) to allow for exploration for 5 minutes. On training day (day 2), the animal started in the light side, the door opened, and they crossed to the dark side, where they received one mild shock (0.5 mA) and then were removed. On the testing days (days 3-5), animals started in the light side and latency to cross to the dark side was measured automatically by the software. The animals are removed after 10 minutes if they do not cross.

#### Three Chamber Social Threat

Social threat was assessed using a modified 3 chamber social interaction task. In this case, the interactors are retired Long Evans breeders, which represent an aggressive threat. The test consisted of 5 phases: habituation, training, and 3 days of testing. On habituation day (day 1), experimental animals were placed in the center chamber and free to explore the three-chamber apparatus for 10 minutes. The two outer chambers contained the empty interactor carousels. On training day (day 2), an aggressive Long Evans was placed in one of the carousels in the left or right chamber with a checkered wall providing a contextual cue in the aggressor’s chamber. The experimental animal was restricted to in that chamber for the first 5 minutes. The door was then removed, and the experimental animal was free to leave that chamber and move freely to all chambers for 5 more minutes. On testing days (days 3-5), the carousels were empty with the checkered wall context around the aggressor’s chamber from the training day, and the experimental animal was placed in the center chamber and allowed to freely to explore for 10 minutes. At all stages, experimental animals were tracked using Ethovision 12 (Noldus) to determine the time they spent in each chamber.

#### Acoustic Startle

Acoustic startle was run using the Gemini system (San Diego Instruments). This one-day test was conducted in an insulated chamber where the animals were lightly restrained in a tube that measures startle response to auditory tones. Variable tones (0, 95, 110, 120 dB) were presented at random intervals for 24 trials, with each tone presenting 6 times. Average and max startle responses (mV) was calculated for each tone.

#### Fear Conditioning

Cued fear conditioning was run using Ethovision 12 (Noldus) and consisted of 5 phases: acquisition, 3 days of extinction, and reinstatement. Acquisition (day 1) consisted of 6 shock-tone pairings (shock intensity was 0.4 mA), each extinction day (days 2-4) consisted of 7 tone presentations, and reinstatement (day 7) consisted of 5 shock-tone pairings. Percent freezing was scored automatically each day using Ethovision’s tracking. Freezing was determined for the total trial, as well as specifically during tones and intertrial intervals (ITIs).

#### Statistics

One-way ANOVAs and Two-way repeated measures ANOVAs (GraphPad Prism 9) were primarily used to assess behavioral endpoints. Outliers were determined by values that fall outside the mean ± 1.96 times the standard deviation. For passive avoidance one-way ANOVAs were used to assess crossing latency on each day. For social threat, Group x Day ANOVAs were used to assess time in the “threatening” side. For acoustic startle, one-way ANOVAs were used to assess startle responses to each of the variable tones. For fear conditioning, Group x Trial ANOVAs were used to analyze freezing during tones and ITIs on individual days, and a Group x Day ANOVA was used to analyze average freezing across the phases. All post hoc testing utilized Tukey tests.

### Fos Activation in the BLA Following Fear Conditioning

In order to tie these behaviors back to the BLA, we analyzed Fos activation 90 minutes following reinstatement. Brains were collected and stored as described in Experiment 4.

#### Immunohistochemistry

A triple label for WFA/PV/Fos was run using the above WFA and PV markers and a new Fos antibody (Santa Cruz, rabbit anti-Fos, sc-253, 1:1000). Fluorescence IHC was run using the same protocol described in Experiment 4.

#### Image Analysis and Quantification

Images were obtained using a Nikon C1 confocal microscope (Nikon Instruments Inc). Z-stacks (0.4 μm steps) were captured using the 20x objective. Images were taken bilaterally from 3 levels of the BLA (approx. at AP -2.0, -2.5, and -3.0) and analyzed using Image J. Image settings were kept constant for all animals. All counts were collected by a blinded observer. BLA cells were determined to be positive or negative for WFA, PV, and Fos, yielding counts for all combinations. One and two-way ANOVAs, followed by Tukey’s post hocs, were used where appropriate.

### Experiment 6: Testing the Necessity of BLA PNNs in Loss-like Behaviors

While the above evidence suggests that the increase in BLA PNNs following ER plays an important role in the development of loss-like behavioral phenotypes, they do not establish a solid causal link between the two. As such, we sought to experimentally probe this relationship using Chondroitinase ABC (ChABC), a plant enzyme which digests ECM and PNNs. By digesting PNNs during the removal period, we can test the necessity of the accumulation of PNNs in loss-like behaviors.

#### ChABC Pilot

ChABC enzymatically digests PNNs rapidly following injection, and then PNNs gradually repopulate over the next few months^23,33^. This creates a period in which we can use ChABC to test the necessity of PNNs during removal. In order to establish the degree and timing of PNN depletion in our hands, we ran a pilot in which rats received unilateral BLA ChABC injections (Sigma, Chondroitinase ABC from Proteus vulgaris, C3667). Two doses (50 U/ml and 200 U/ml ChABC in 1xPBS) and volumes (500 nl and 1000 nl) were tested (n=6/group), and injections were counterbalanced for side. Half of the rats were sacrificed 4-5 days after surgery to determine the initial degree of PNN depletion, and the other half were sacrificed 3 weeks after surgery to determine the degree of PNN repopulation following ChABC injection at that time. Brains were collected and WFA/PV IHCs were run as described above. WFA area in the BLA was measured bilaterally, enabling comparison between the side that received a ChABC injection and the uninjected side. Paired t-tests were utilized in this analysis. Altogether, ChABC resulted in a ∼90% depletion at 4-5 days and remained at a ∼60% depletion 3 weeks later (Figure 7B). A dose of 200 U/ml and volume of 500 nl were determined to be the optimum parameters for the experiment.

#### Experimental Design

This experiment utilized a 2×2 design, with removal (EE vs ER) and treatment (VEH vs ChABC) as factors (n=12/group) (Fig 7A). This study did not include SH groups to allow for the other groups to have a sufficient N, and as EE is the more appropriate control. EE and ER were conducted as described above. At the time of removal, all rats received bilateral injections of either ChABC or VEH into the BLA (500 nl/side of 200U/ml ChABC or 1xPBS; coordinates AP - 2.6, ML +/-5.0, DV -8.0). All were single housed for a 2-day recovery period following surgery, with EE rats being placed back into the active-cycle enrichment paradigm afterwards. No issues were observed following the return to group housing. ER rats remained single housed for the duration of the experiment. This study was run in 2 cohorts each with equal representation from each group. Behavioral testing began 9-12 days following surgery and was concluded by 20-23 days after surgery, in order to remain within the window of PNN depletion established in the pilot. Bodyweight was monitored throughout this period.

Behaviors consisted of passive avoidance (PA), acoustic startle (AS), and forced swim (FST), with a 2-day rest period between each. PA and AS were run and analyzed as described above (Experiment 5). FST was run as a 2-day test, with a 15-minute swim on Day 1 and a 6-minute swim on Day 2. Rats were then sacrificed 90-minutes after the Day 2 swim. Swimming, immobility, climbing, and diving behaviors were hand-scored by a blind observer for the first 6 minutes of each test day^45^. One-way and Two-way ANOVAs, followed by Tukey’s post hocs, were used to analyze each endpoint, where appropriate. Brains were collected to verify hits and misses and to quantify PNN levels in each group at the conclusion of the experiment.

## Supporting information

Supplementary Information Key and Legends

## Funding

This work was supported by MH049698 to JPH, MH107487 and MH121102 to REM, and MH125541 to MAS.

## Author contributions

MAS, JPH, and REM conceptualized the study together. BLS generated animals for Experiments 1 and 2. RS, HME, and MAS analyzed the RNAseq data. JKR and MAS analyzed the proteomics data. KA and MAS ran and analyzed the kinomics data. JLB and MAS analyzed the IHC data. RKP and MAS analyzed the synaptic input data. RDM conceptualized the social threat assay and assisted with design of behavioral analyses. HP, EW, and NN participated in interpreting the results. MAS generated figures and wrote the manuscript. MAS, JPH, and REM edited the manuscript. All authors participated in finalizing the manuscript.

## Acknowledgements

The authors would like to thank Ali S Imami and Xiaolu Zhang for consulting on the RNAseq analysis; Adam Funk for consulting on the proteomics analysis; Evelin Cotella for consulting on the behavioral tests; and Parinaz Mahbod and Sinead O’ Donovan for consulting on tissue processing.

## Data and materials availability

RNAseq data are available at Gene Expression Omnibus under accession number GSE208029. All other data are available in the main text or supplementary materials.

## Conflict of interest

The authors declare no competing interests.

## Notes

### Competing Interest Statement

The authors have declared no competing interest.

### Summary of Updates

An additional experiment has been added to the manuscript to test the necessity of BLA ECM accumulation in loss-like behaviors.

