## Supplementary Information Key and Legends for "Molecular Neurobiology of Loss"

**Table of Contents**

**Extended Data Figures** (*each as separate PDF file*):

- **Extended Data 1**: Bodyweights for Experiments 1-5
- **Extended Data 2**: FOS Screen Results
- **Extended Data 3**: Overview of Bioinformatics Analysis Pipeline
- **Extended Data 4**: Full Results for Experiment 2 RNAseq
- **Extended Data 5**: Full Results for Experiment 3 RNAseq
- **Extended Data 6**: Full Results for Experiment 3 Proteomics and Kinomics
- **Extended Data 7**: Additional IHC Results
- **Extended Data 8**: Additional Behavior Results from Experiment 5
- **Extended Data 9**: Additional Behavior Results from Experiment 6

**Supplementary Tables** (*each as separate Excel file*):

- **Table S1**: Overview of Experiments
- **Table S2**: RNAseq Gene Expression Data
- **Table S3**: Proteomics and Kinomics Data
- **Table S4**: Pathway Analysis Results
- **Table S5**: Perturbagen Analysis Results
- **Table S6**: Cell Type Analysis Results
- **Table S7**: Microglia and ECM Focused Gene Expression
- **Table S8**: Detailed Statistical Results

**EXTENDED DATA FIGURES** *(each as separate PDF file)*

**Extended Data 1: Bodyweights for Experiments 1-5.** ER consistently causes rats to gain weight (A-E). Data are presented as bodyweight percent change from weight at the time of enrichment removal. Note x-axis as days post removal, as visual differences here are due to differences in weight sampling times, not differences between experiments. In all experiments, ER rats gained more weight than EE and SH rats. * = p<0.05.

**Extended Data 2: FOS Screen Results.** SH, EE, and ER rats were sacrificed 120 minutes after the forced swim test. IHC was used to explore the FOS response in various stress-responsive brain regions. Representative images are shown for the regions that showed the most differential response between groups. * = p<0.05. IL = infralimbic prefrontal cortex, MEA = medial amygdala. PL = prelimbic prefrontal cortex. CEA = central amygdala. BST = bed nucleus of the stria terminalis. PVN = paraventricular nucleus. NAc = nucleus accumbens. DG = dentate gyrus.

**Extended Data 3: Overview of Bioinformatics Analysis Pipeline.** Multi-Omics analyses were conducted as depicted from left to right, with data inputs, bioinformatics tools used, data condensing steps, and endpoints shown where appropriate. The ultimate goal of these analyses was to identify commonly implicated processes, pathways, and targets, with the notion that cross-analysis and cross-platform validation will yield higher-confidence results.

**Extended Data 4: Full Results for Experiment 2 RNAseq.** (A) Full transcriptome pathway analysis results. Significant pathways (p<0.05) from GSEA were condensed based on semantic similarity using Pathway Hunter (small left labels, and these categories were further condensed into functional themes based on *a priori* knowledge (leftmost labels). Enrichment scores for individual pathways are represented by the heatmap (with EEvSH on the left, ERvSH in the middle, and ERvEE on the right), and the degree of each category’s enrichment is represented by the p value dots on the right. The enrichment scores show how much (magnitude) and in what direction (yellow is up, blue is down) a theme was changed by the model, while the p values show the contribution of each category to these changes. (B) Targeted pathway analysis results. Analysis is the same as described in A but uses significant pathways (p<0.05) from Enrichr. (C) Themes that replicated between these two pathway analyses are shown in the replicated pathways heatmap, with darker green indicating more involvement. Right labels are Pathway Hunter categories, and left labels are their larger themes. (D) Signature analysis results. iLINCS perturbagen signatures that are concordant with EEvSH or discordant with ERvSH and ERvEE would theoretically reverse ER, while signatures that are discordant with EEvSH or concordant with ERvSH and ERvEE would theoretically mimic ER. MOAs (small left labels) for the top 20 perturbagens were organized into functional themes (far left label). Purple represents the number of perturbagens with that MOA in that contrast. (E) Cell type analysis results.

**Extended Data 5: Full Results for Experiment 3 RNAseq.** Data is presented as described in Extended Data 4. (A) Full transcriptome pathway analysis results. (B) Targeted pathway analysis results. (C) Replicated pathways. (D) Signature analysis results. (E) Cell type analysis results.

**Extended Data 6: Full Results for Experiment 3 Proteomics and Kinomics.** (A) Proteomics pathway results. Pathways significantly enriched (p<0.05) in the top 100 upregulated and top 100 downregulated peptides. Data is presented as described in Extended Data 4A. (B) Kinomics pathway results. Pathways significantly enriched (p<0.05) in the differentially phosphorylated (log2FC>0.3) peptides. Data is presented as described in Extended Data 4A. (C) Upstream kinases identified by KRSA. Z-scores are presented for each kinase in each group, with higher z-scores indicating stronger over- (yellow) or under- (blue) representation. (D) Replicated pathways across Experiment 3 RNAseq, proteomics, and kinomics. Data is presented as described in Extended Data 4C.

**Extended Data 7: Additional IHC Results.** Other BLA endpoints from Experiments 4 and 5. (A) Microglia counts. (B) Microglia intensity. (C) Microglia soma size. (D) Parvalbumin counts. (E) WFA intensity. No differences were observed in these measures. (F) vGAT puncta counts separated by group. (G) vGLUT puncta counts separated by group. Note that PNN phenotypes were observed in all conditions. * = p<0.05

**Extended Data 8: Additional Behavior Results from Experiment 5.** Other behavioral endpoints from Experiments 5. (A) Passive avoidance latency to cross on habituation, training, and testing days. ER rats exhibited a moderately faster latency to cross, but this effect did not emerge until testing day 3. (B) Additional social threat endpoints, including time in each zone on habituation and training days, time to leave the retired breeder long evans (LE) zone on training day, and overall locomotion. All groups showed a bias for the left compartment that persisted across days. The presence of the LE caused EE animals to spend less time in this zone. No differences were observed in time to leave the LE zone. ER rats did travel a greater distance than EE rats on all testing days, possibly pointing to increased escape behavior. (C) Average and maximum startle intensity across tones. ER rats responded more to the higher tones for both endpoints. (D) Fear Conditioning extinction day 2 and 3 freezing during ITIs. ER and SH rats showed enhanced extinction compared to EE rats. Freezing during fear conditioning tones each day. Patterns are similar to those seen during ITIs (Figure 5E), but of a smaller magnitude. * = p<0.05

**Extended Data 9: Additional Behavior Results from Experiment 6.** Other behavioral endpoints from Experiments 6. (E) Passive avoidance latency to cross on habituation, training, and testing days 2 and 3. Other testing days exhibited the same pattern observed on testing day 1 (Figure 7D). ER VEH rats showing an increased latency to cross, with ChABC blocking this effect. (D) Average and maximum startle intensity across tones. ER rats responded more to the highest tone for both endpoints. (F) Additional forced swim test results. ER VEH rats showed an increased latency to immobility and decreased swimming. ChABC blocked both effects in ER rats. No differences were observed for climbing and diving behaviors. The same patterns were observed on Day 2, with a slight shift toward increased mobility observed across all groups. * = p<0.05

**SUPPLEMENTARY TABLES** *(each as separate Excel file)*

**Table S1: Overview of Experiments.** This study consisted of 6 experiments. The goals and approaches of each experiment are detailed here.

**Table S2**: **RNAseq Gene Expression Data.**

*Tab 1:* Experiment 2 *Differential Gene Expression*. Experiment 2 RNAseq differential expression values for EEvSH, ERvSH, and ERvEE genes. Gene names are provided in column A, and differential expression values are presented in log2FC.

*Tab 2:* Experiment 3 *Differential Gene Expression*. Experiment 3 RNAseq differential expression values for EEvSH, ERvSH, and ERvEE genes. Gene names are provided in column A, and differential expression values are presented in log2FC.

**Table S3: Proteomics and Kinomics Data.**

*Tab 1: Proteomics Differential Expression.* Differential expression of peptides detected via LCMS, presented in log2FC.

*Tab 2: Kinomics Differential Phosphorylation.* Differential phosphorylation of STK peptides, presented in FC.

*Tab 3: Kinomics KRSA.* Upstream kinase analysis. Z-score signifies how over- (positive) or under- (negative) represented a particular kinase’s activity is in that contrast. Kinomics was run in triplicate on 3 separate STK chips, so z-scores are provided for each chip and as an overall average.

*Tab 4: Kinomics KRSA Summary*. Summary of average z-scores from Tab 3 for kinases considered to be strongly represented (>2.0), represented (1.75-2), or weakly represented (1.5-1.75). Kinases with z-scores <1.5 are not considered to be differentially active in the sample. Corresponds to Extended Data 6C.

**Table S4: Pathway Analysis Results**

*Tab 1: Experiment 2 GSEA Pathways*. Detailed pathways for Experiment 2’s RNAseq Full Transcriptome Pathway Analysis. Corresponds to Figure 2A and Extended Data4A. IDs and titles of Individual pathways are provided in columns D and E, respectively. Enrichment Scores for that pathway in each contrast are provided in columns F-H, with yellow signifying positive scores, blue signifying negative scores, and while signifying that that pathway was not significantly enriched in that contrast. Pathways are organized into clusters based on semantic similarity by PathwayHunter. PathwayHunter cluster names, numbers, and p values (how enriched that cluster is overall) are provided in columns B, C, and I. The larger themes that these categories were organized into based on *a priori* knowledge are provided in column A. This file can be used to look up which pathways make up the categories and themes presented in the main body of the manuscript.

*Tab 2: Experiment 2 Enrichr Pathways.* Detailed pathways for Experiment 2’s RNAseq Targeted Pathway Analysis. Corresponds to Extended Data 4B. See Tab 1 legend for key. Individual pathway enrichments are presented as Enrichr Combined Scores, rather than GSEA Enrichment Scores.

*Tab 3: Experiment 2 Replicated Pathways.* Themes of pathways observed in both GSEA and Enrichr results. Numbers correspond to -log10pvalue enrichment of PathwayHunter categories (on right). Overarching themes provided on left. Corresponds to Extended Data 4C.

*Tab 4*: *Experiment 3 GSEA Pathways.* Detailed pathways for Experiment 3’s RNAseq Full Transcriptome Pathway Analysis. Corresponds to Extended Data 5A. See Tab 1 legend for key.

*Tab 5: Experiment 3 Enrichr Pathways.* Detailed pathways for Experiment 2’s RNAseq Targeted Pathway Analysis. Corresponds to Extended Data 5B. See Tab 2 legend for key.

*Tab 6*: Experiment 3 Replicated Pathways. Themes of pathways observed in both GSEA and Enrichr results. See Tab 3 legend for key. Corresponds to Extended Data 5C.

*Tab 7: Experiment 3 Proteomics Pathways.* Detailed pathways for Experiment 2’s Proteomics Pathway Analysis. Corresponds to Extended Data 6A. See Tab 2 legend for key.

*Tab 8: Experiment 3 Kinomics Pathways.* Detailed pathways for Experiment 2’s Kinomics Pathway Analysis. Corresponds to Extended Data 6B. See Tab 2 legend for key.

*Tab 9: Experiment 3 All Replicated Pathways.* Themes of pathways observed in RNAseq, proteomics, and kinomics results. See Tab 3 legend for key. Corresponds to Extended Data 6D.

*Tab 10: Overall Replicated Pathways.* Themes of pathways observed in Experiment 2 RNAseq results and Experiment 3 RNAseq, proteomics, and kinomics results. Corresponds to Figure 2B. See Tab 3 legend for key.

**Table S5: Perturbagen Analysis Results**

*Tab 1: Experiment 2 Perturbagens*. Top 20 concordant (+) and discordant (-) perturbagens for EEvSH, ERvSH, and ERvEE. ID, name, and mechanism of action are given for each perturbagen, along with the p value and z-score representing the significance and strength of similarity between that perturbagen’s L1000 signature and the experimental L1000 signature.

*Tab 2: Experiment 2 Summary.* Summary of MOAs expected to reverse or mimic ER signatures. Numbers correspond to the number of perturbagens that had that MOA for each contrast, calculated from the individual perturbagens in Tab 1. Corresponds to Extended Data 4D.

*Tab 3: Experiment 3 Perturbagens*. Top 20 concordant (+) and discordant (-) perturbagens for EEvSH, ERvSH, and ERvEE. ID, name, and mechanism of action are given for each perturbagen, along with the p value and z-score representing the significance and strength of similarity between that perturbagen’s L1000 signature and the experimental L1000 signature.

*Tab 4: Experiment 3 Summary.* Summary of MOAs expected to reverse or mimic ER signatures. Numbers correspond to the number of perturbagens that had that MOA for each contrast, calculated from the individual perturbagens in Tab 3. Corresponds to Extended Data 5D.

**Table S6: Cell Type Analysis Results**

*Tab 1: Experiment 2 Leading Edge Genes.* Top 100 upregulated and top 100 downregulated leading-edge genes from GSEA are provided for EEvSH, ERvSH, and ERvEE in columns B, L, and V, respectively. Expected expression of these genes in different cell types, named in row 2, are given next to each gene.

*Tab 2: Experiment 2 Average Cell Types.* Average leading-edge gene expression in different cell types, calculated from the individual gene values in Tab 1. Corresponds to Figure 3A and Extended Data 4E.

*Tab 3: Experiment 3 Leading Edge Genes.* Top 100 upregulated and top 100 downregulated leading-edge genes from GSEA are provided for EEvSH, ERvSH, and ERvEE in columns B, L, and V, respectively. Expected expression of these genes in different cell types, named in row 2, are given next to each gene.

*Tab 4: Experiment 3 Average Cell Types.* Average leading-edge gene expression in different cell types calculated from the individual gene values in Tab 3. Corresponds to Extended Data 5E.

**Table S7: Microglia and ECM Focused Gene Expression.** Differential gene expression related to microglia and ECM. Corresponds to Figure 2C.

**Table S8: Detailed Statistical Results.**

*Tab 1: Figure 1*. Statistical results for Figure 1B.

*Tab 2: Figure 3*. Statistical results for Figure 3A-C.

*Tab 3: Figure 4*. Statistical results for Figure 4B-D, F, G, I-K

*Tab 4: Figure 5*. Statistical results for Figure 5B-C, E.

*Tab 5: Figure 6*. Statistical results for Figure 6B-D.

*Tab 6: Figure 7*. Statistical results for Figure 7B-F.

*Tab 7: Extended Data 1*. Statistical results for Extended Data 1A-E.

*Tab 8: Extended Data 2*. Statistical results for Extended Data 2.

*Tab 9: Extended Data 7*. Statistical results for Extended Data 7A-G.

*Tab 10: Extended Data 8*. Statistical results for Extended Data 8A-D.

*Tab 11: Extended Data 9*. Statistical results for Extended Data 9A-C.
