## Supplementary figures and images for "Molecular Neurobiology of Loss"

### Extended Data 1

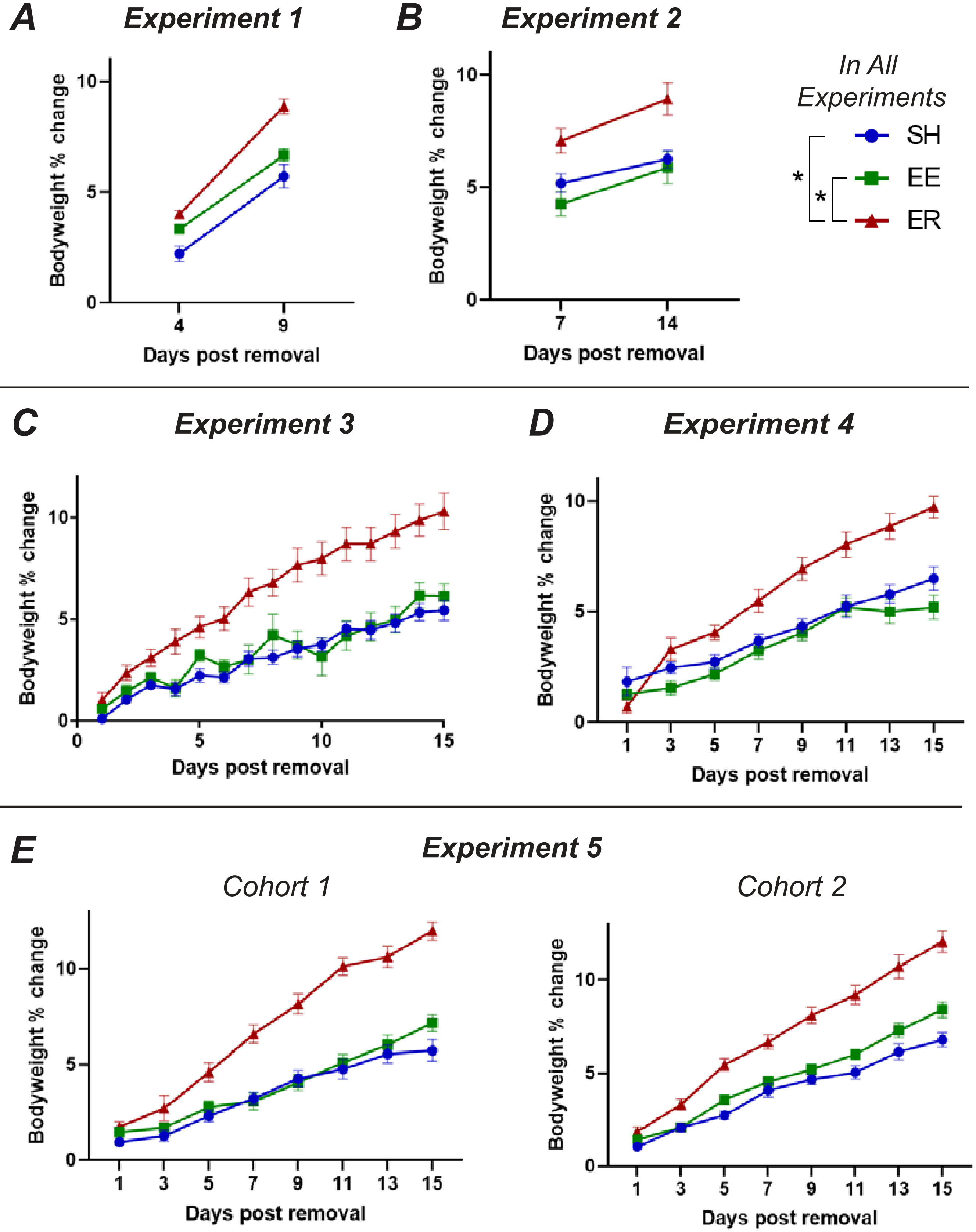

### Extended Data 2

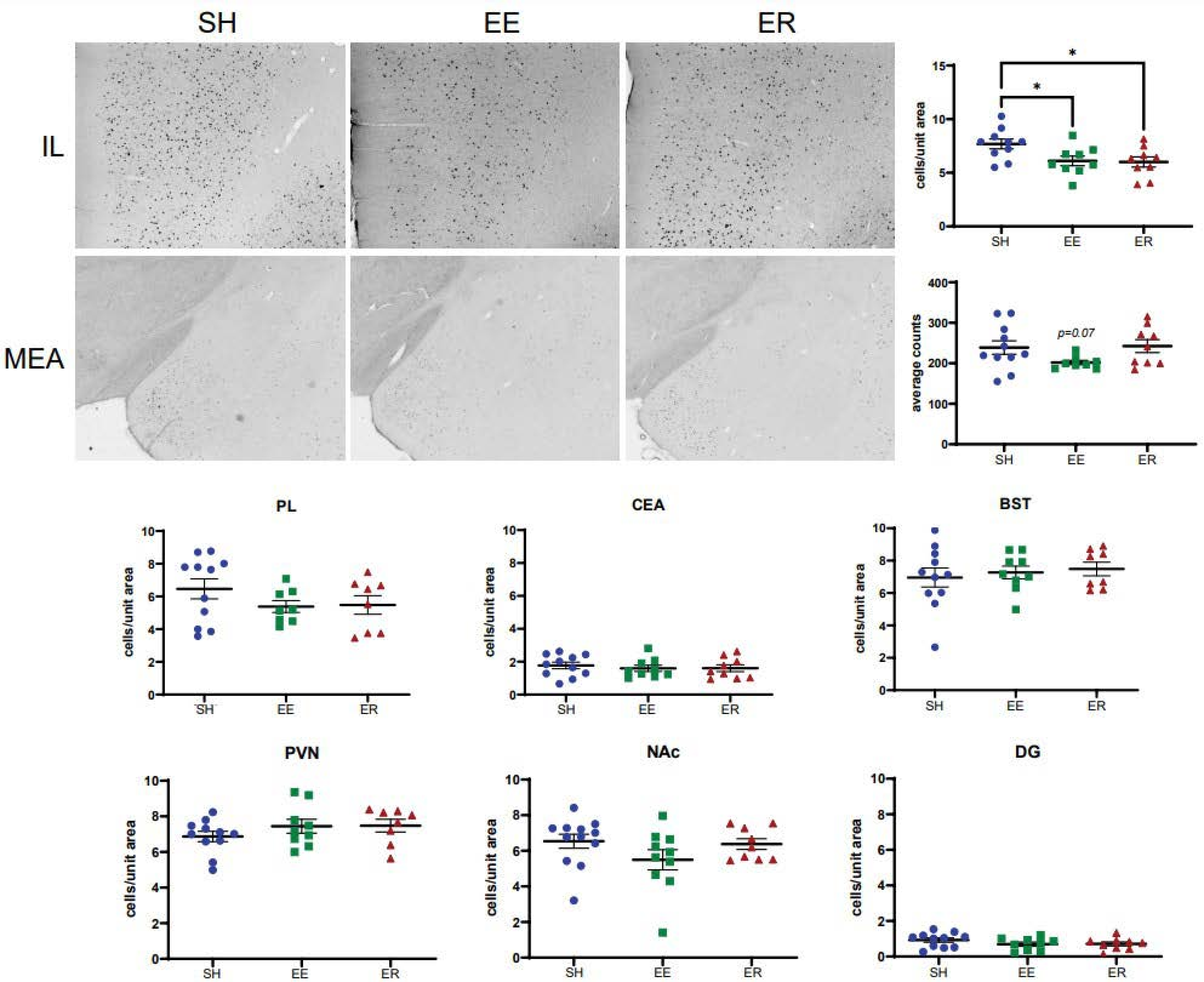

### Extended Data 3

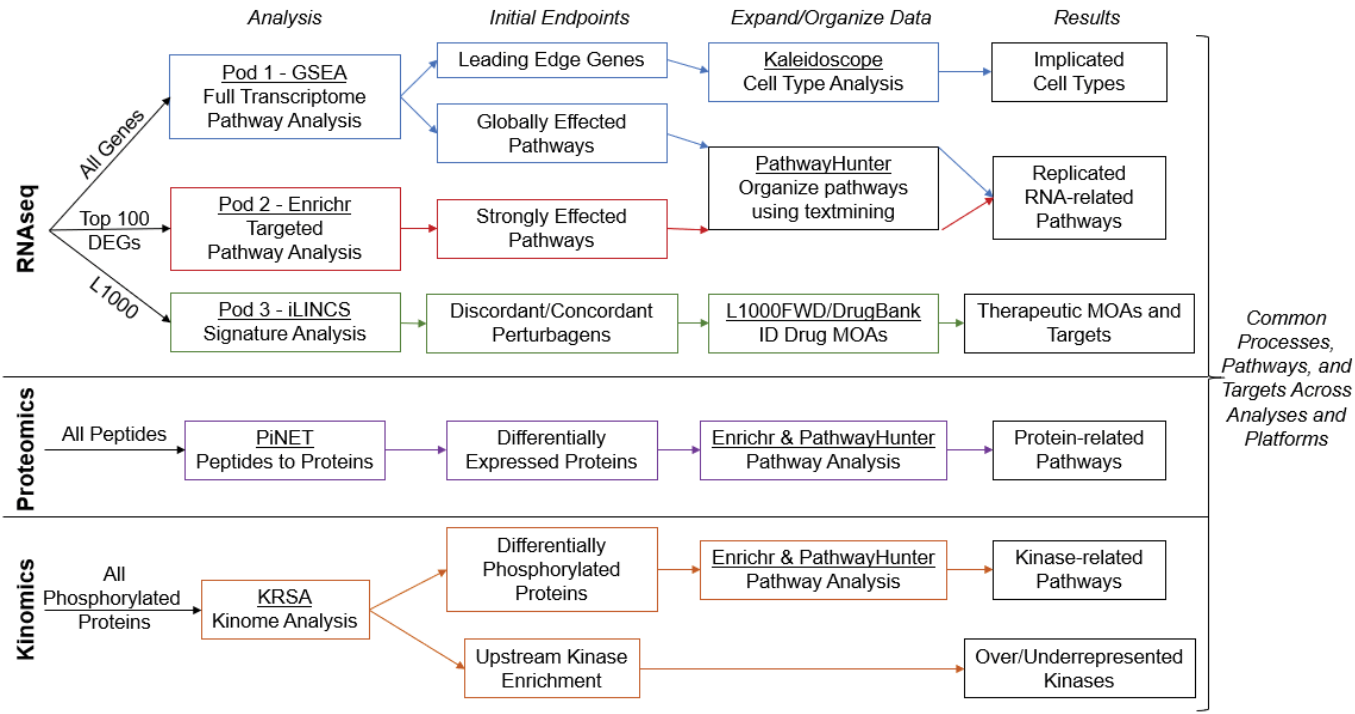

### Extended Data 4

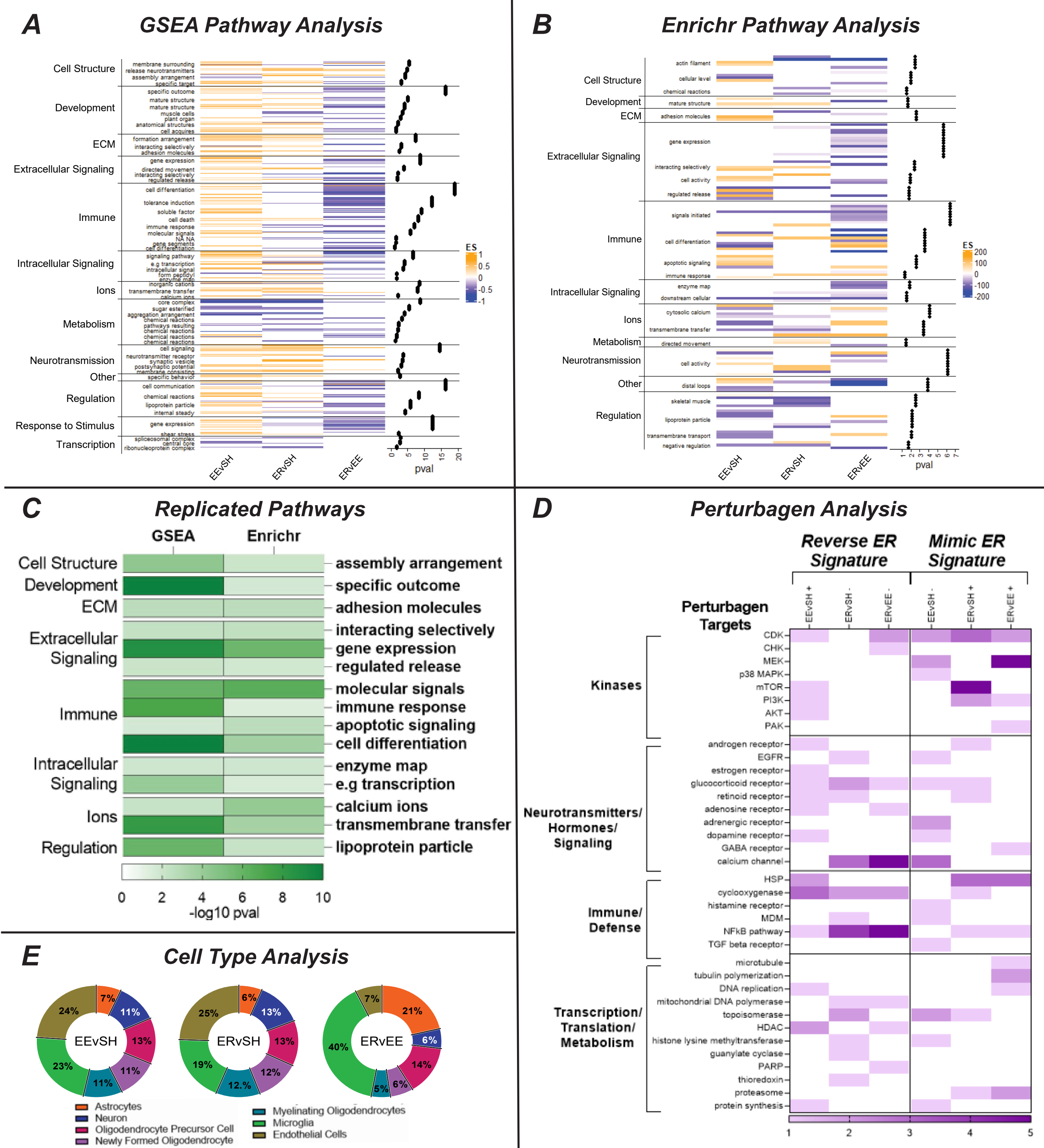

### Extended Data 5

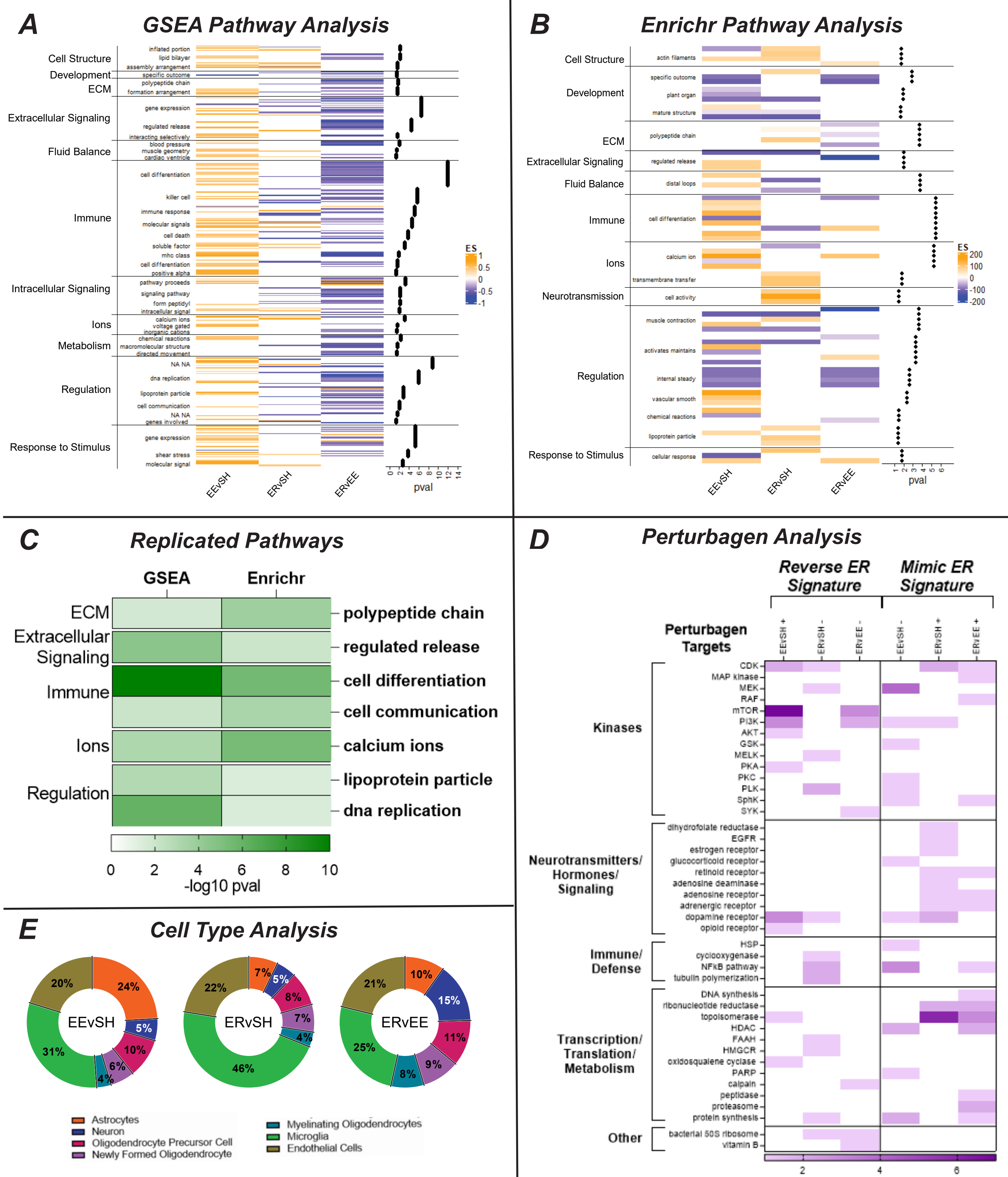

### Extended Data 6

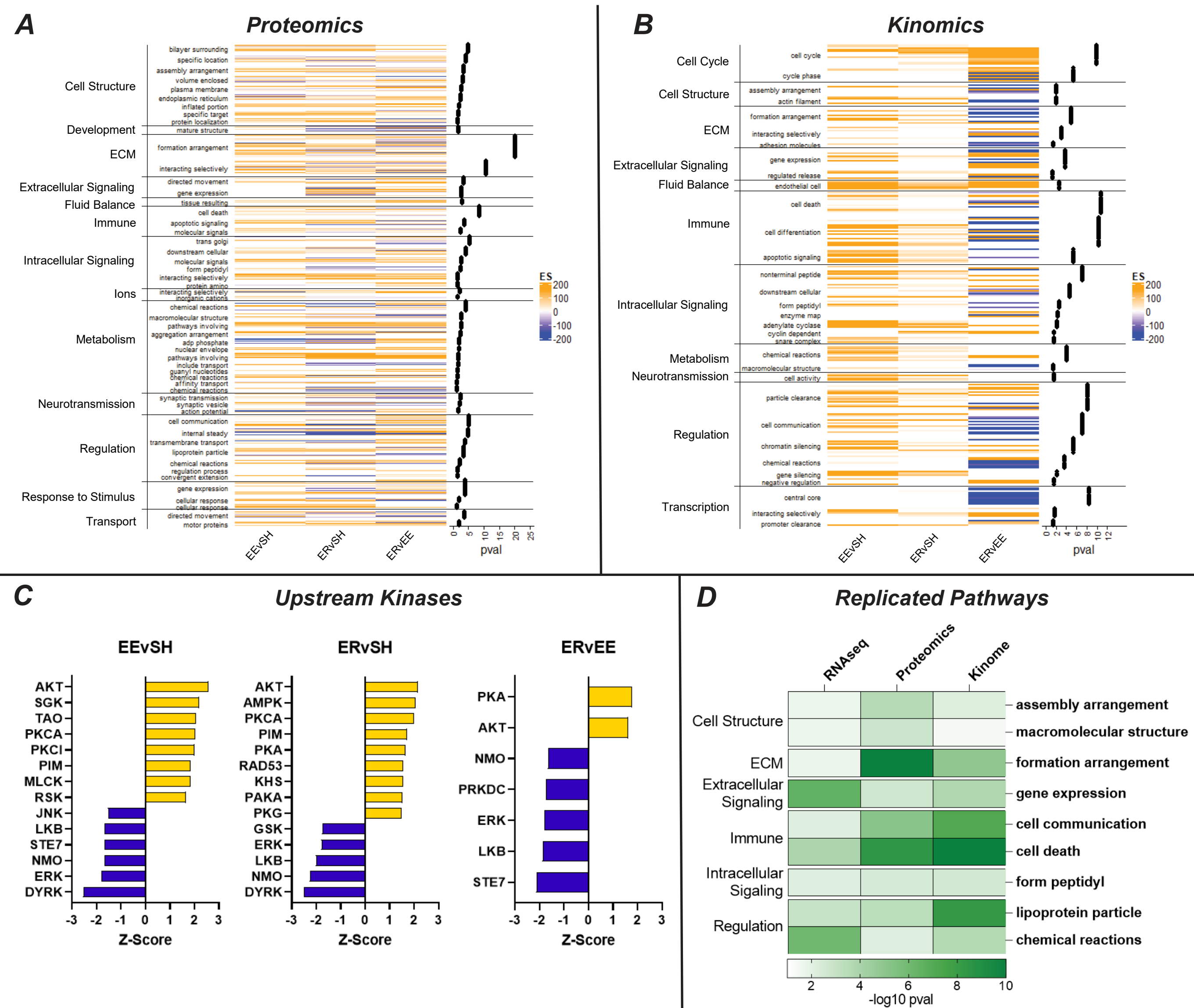

### Extended Data 7

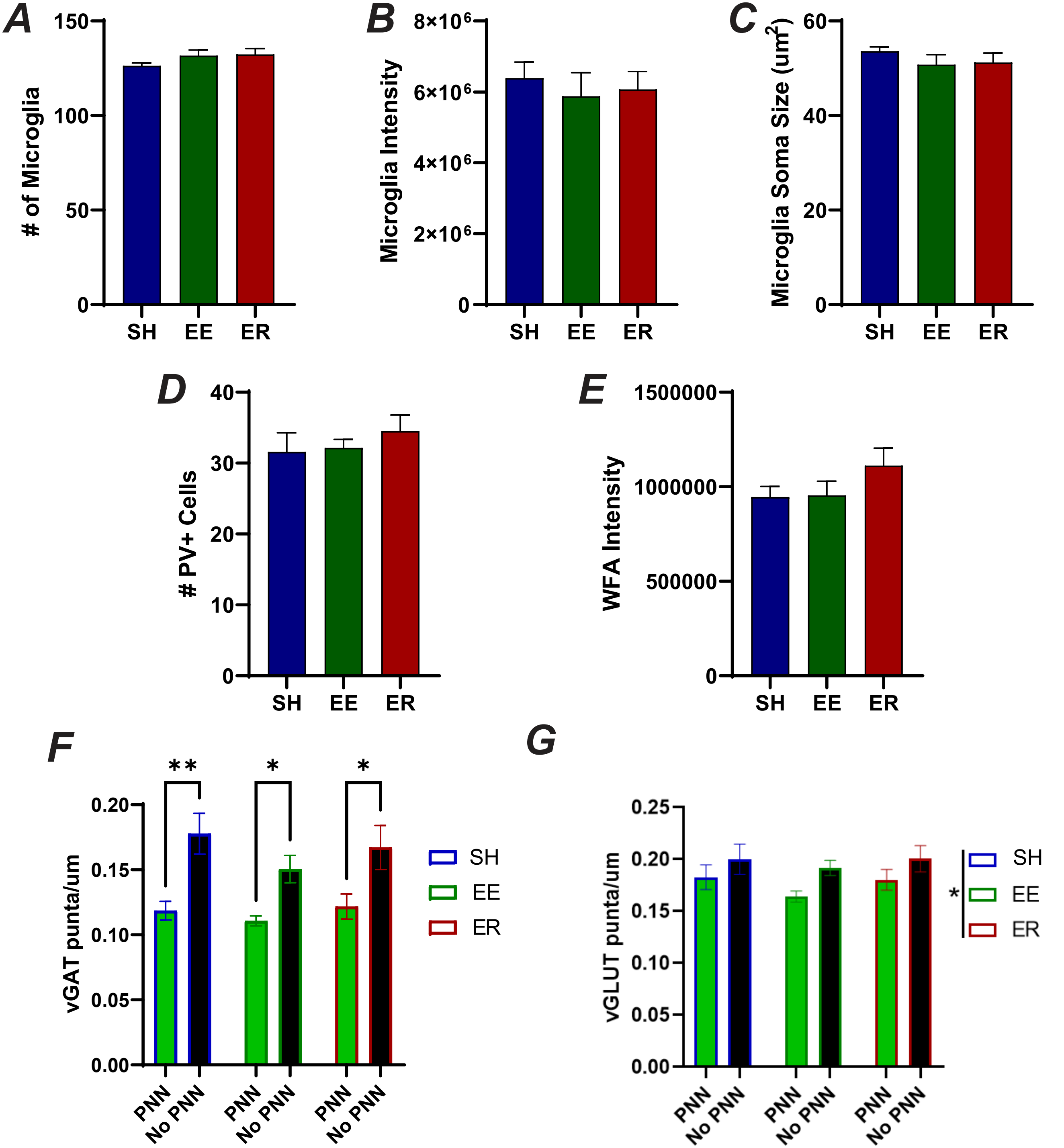

### Extended Data 8

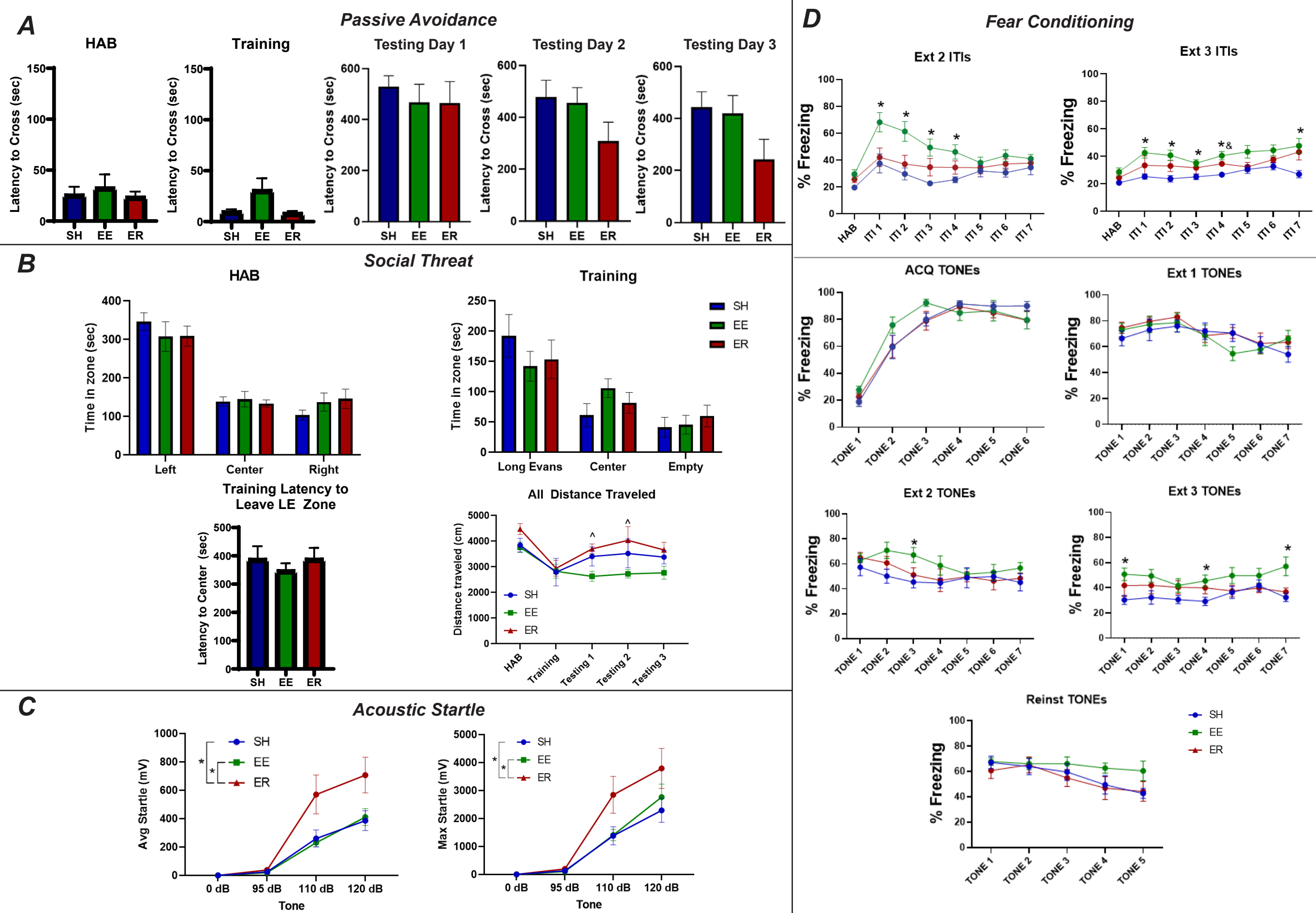

### Extended Data 9

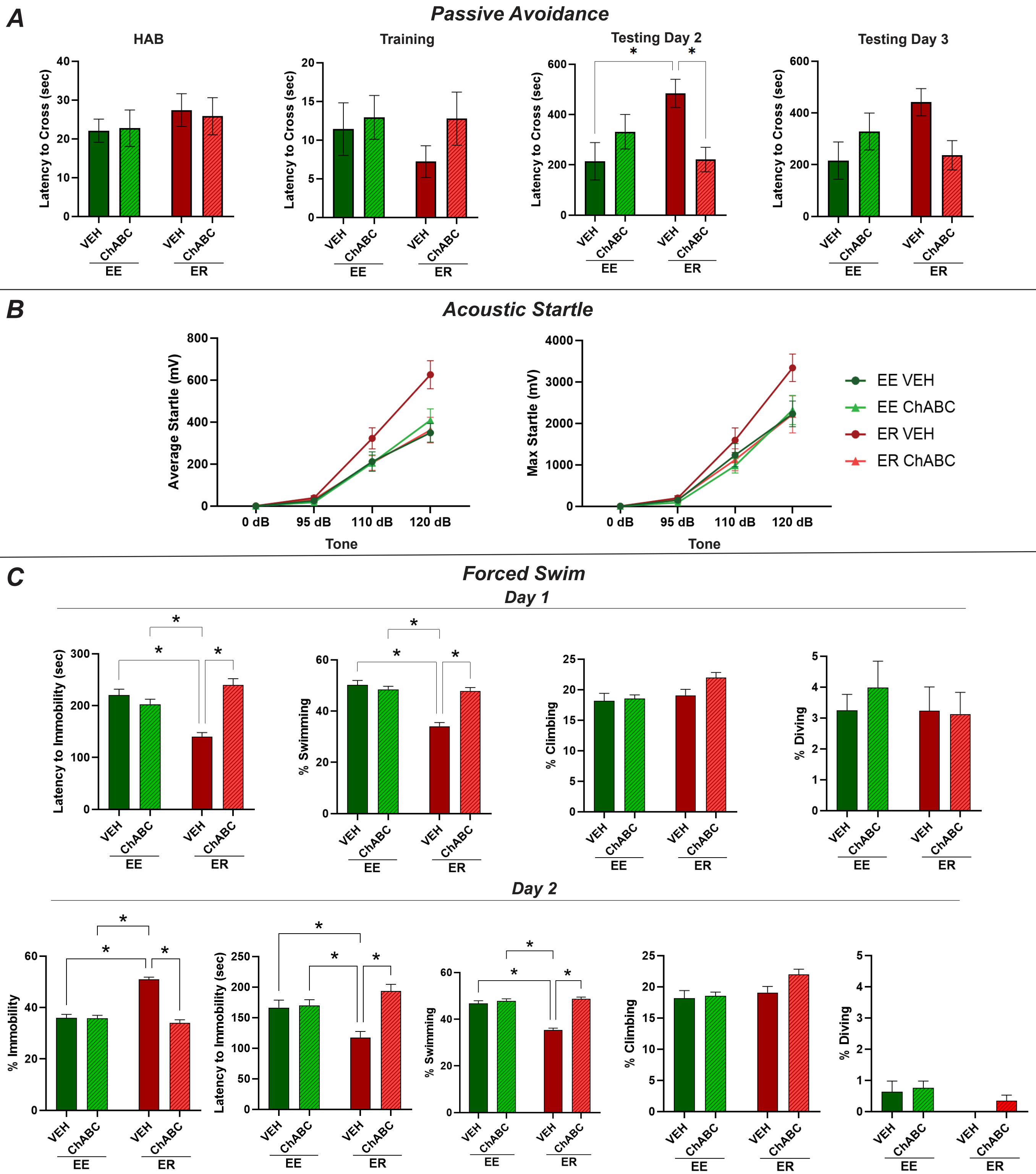
